# Evaluating regional heritability mapping methods for identifying QTLs in a wild population of Soay sheep

**DOI:** 10.1101/2024.06.08.598050

**Authors:** Caelinn James, Josephine M. Pemberton, Pau Navarro, Sara Knott

## Abstract

Regional heritability mapping (RHM) is a method that estimates the heritability of genomic segments that may contain both common and rare variants affecting a complex trait. We compared three RHM methods: SNP-RHM, which uses genomic relationship matrices (GRMs) based on SNP genotypes; Hap-RHM, which uses GRMs based on haplotypes; and SNHap-RHM, which uses both SNP-based and haplotype-based GRMs jointly. We applied these methods to data from a wild population of sheep, analysed eleven polygenic morphometric traits and compared the results with previous genome wide association analyses (GWAS). We found that whilst the inclusion of the regional matrix did not explain significant variation for all regions that were associated with trait variation using GWAS, it did for several regions that were not previously associated with trait variation.

## Introduction

Genome-wide association studies (GWAS) are commonly used to identify genotyped SNPs in linkage disequilibrium (LD) with causal loci. The regions around the SNPs associated with the focal trait can then be examined as these SNPs serve as markers for the causal loci. For example, the function of nearby genes can be investigated to see if they are involved in biological pathways related to the trait, or fine-mapping can be performed to narrow down the relevant region and pinpoint the causal variant. However, GWAS has some limitations and challenges that prevent it from finding all the genetic factors that contribute to complex traits. One of these limitations is the power of GWAS, which is the ability to detect true associations. The power of GWAS depends on several factors, such as the sample size, the variant effect size, whether genotyped SNPs are in LD with causal SNPs, and the allele frequency of the causal variant.

To overcome the limitations of GWAS, especially when a trait is influenced by multiple independent effects and/or rare variants in a region, regional heritability mapping (RHM) methods have been developed (Nagamine et al. 2012; Shirali et al. 2018; Oppong et al. 2021). RHM is a technique that estimates the heritability of a trait that is explained by a specific region of the genome. To estimate the heritability of a region, RHM uses a genomic relationship matrix (GRM), which is a matrix that captures the genetic similarity between individuals based on their SNP genotypes in that region. RHM also corrects for the genetic similarity across the whole genome by fitting another GRM that includes all the SNPs in the genome (or a leave-one-chromosome-out (LOCO) GRM that excludes the chromosome where the region of interest is located). By comparing the model fit with and without the regional GRM (rGRM), RHM can identify regions that contain causal variants for the trait, and by using the variance estimate for the rGRM, RHM can estimate how much heritability that region contributes.

RHM can be performed using different types of rGRMs and region sizes, depending on the assumptions and goals of the analysis. There are three main types of RHM that have been proposed. The first type is SNP-RHM, which uses rGRMs that are based on the sharing of SNP alleles across a region. The regions are usually defined as windows that contain a fixed number of SNPs (Nagamine et al. 2012). SNP-RHM aims to identify regions with multiple SNPs that are in LD with the multiple causal variants that have too small an effect on the trait individually to be detected by GWAS. However, SNP-RHM only captures effects associated with genotyped SNPs. The second type is Hap-RHM, which uses rGRMs that are based on the sharing of haplotype alleles across a region. The regions are defined as haplotype blocks (Shirali et al. 2018). Hap-RHM aims to identify regions where the causal variant is in LD with the haplotype allele, but not necessarily with any specific genotyped SNPs, which allows for detection of variance that is not captured by genotyped SNPs. This method can capture the effect of rare causal variants due to rare haplotype alleles being more likely to be in LD with rare variants than individual, genotyped SNPs. In addition, haplotype effects may reflect the interaction effects of closely linked causal variants. The third method, SNHap-RHM, simultaneously fits two rGRMs: one SNP-based and one haplotype-based, and defines regions as haplotype blocks (Oppong et al. 2021). This combines the advantages of both SNP-RHM and Hap-RHM to increase power to detect regions containing variants influencing the phenotype. On occasions where SNP- RHM and Hap-RHM can detect genetic variance in the same haplotype block, SNHap-RHM can also be used to give more insight into the underlying genetic architecture.

Here, we evaluate the three RHM methods for their ability to identify regions containing potentially causal loci in a sample of wild Soay sheep. In this study, we analysed 11 polygenic morphometric traits in the Soay sheep population using RHM. These traits include the same traits measured at different ages, as they are affected by different non-genetic factors (and potentially different genetic factors) and vary in heritability across different stages of life. Despite using various methods to search for the genetic variants that affect these traits, such as GWAS (Bérénos et al. 2015; James et al. 2022), genomic prediction (Ashraf et al. 2021) and chromosome partitioning (Bérénos et al. 2015), most of the genetic variation in these traits remains unexplained by the genotyped and imputed SNPs. Moreover, for some of these traits, there are no SNPs that show significant association with the trait variation to date.

Our aims were as follows:

1. To determine the suitability of RHM methods for the Soay sheep data given the smaller sample sizes, lower density SNP data and more potential for missing data in comparison to the human data for which these methods were developed.
2. To compare the results of RHM with those of GWAS to determine the extent to which RHM recovers known associations and identifies new associations.
3. To investigate regions for which including regional matrices in the RHM framework improves model fit, to better characterise the underlying genetic architecture of the focal traits and identify potential causal genes based on known functional data.

## Methods

### Phenotypic data

The Soay sheep (*Ovis aries*) is a primitive breed of sheep that lives on the St. Kilda archipelago. Since 1985, a long-term, individual-based study has been conducted on the population residing on the island of Hirta (Clutton-Brock and Pemberton 2003). Each individual is sampled for DNA analysis and ear-tagged when it is first captured (usually within ten days of birth) so that it can be re-identified later. The study involves regular recaptures to measure various traits throughout an individual’s life, and collection and measurement of skeletal remains after death.

We focused on 11 age-specific morphometric traits which have been repeatedly analysed by different approaches and are known to be polygenic (Bérénos et al. 2015; Ashraf et al. 2021; Hunter et al. 2022; James et al. 2022). We analysed these traits separately by age class (neonate, lamb and adult). Birth weight was the only trait analysed in neonates, defined as individuals who were caught and weighed between two and ten days after birth. In August, lambs (aged approximately 4 months) and adults were caught and measured for weight, foreleg length and hindleg length. Due to adults being recaptured across multiple years, the adult live traits included repeated measurements. Metacarpal length and jaw length were measured from the skeletons after death. We classified “lambs” as individuals who had live trait data recorded in the August of their birth year, or who died before 14 months of age for post mortem measures. We classified “adults” as individuals who had live trait data recorded at least two years after birth, or who died after 26 months of age for post mortem measures. Birth and August weights are recorded to the nearest 0.1kg, whilst the length traits are measured to the nearest mm (Beraldi et al. 2007). We did not analyse yearlings due to low sample size.

See Table 1 for the number of individuals and records per trait.

**Table 1.** Number of individuals and records, fixed and random effects fitted in each trait and age class model during RHM pre-correction, alongside the LOCO GRM.

| Age | Trait | Number of individuals | Number of records | Fixed effects | Random effects |
| --- | --- | --- | --- | --- | --- |
| Neonate | Birth weight | 2975 | 2975 | Sex | Year of birth |
|  |  |  |  | Litter size | Mother ID |
|  |  |  |  | Population size year before birth |  |
|  |  |  |  | Age of mother (quadratic) |  |
|  |  |  |  | Ordinal date of birth |  |
|  |  |  |  | Age (days) |  |
| Lamb | Weight | 2424 | 2424 | Sex | Year of birth |
|  |  |  |  | Litter size | Mother ID |
|  |  |  |  | Population size |  |
|  |  |  |  | Age (days) |  |
|  | Foreleg | 2512 | 2512 | Sex | Year of birth |
|  |  |  |  | Litter size | Mother ID |
|  |  |  |  | Population size |  |
|  |  |  |  | Age (days) |  |
|  | Hindleg | 2577 | 2577 | Sex | Year of birth |
|  |  |  |  | Litter size | Mother ID |
|  |  |  |  | Population size |  |
|  |  |  |  | Age (days) |  |
|  | Metacarpal | 2117 | 2117 | Sex | Year of birth |
|  |  |  |  | Litter size | Mother ID |
|  |  |  |  | Age at death (months) |  |
|  | Jaw | 2172 | 2172 | Sex | Year of birth |
|  |  |  |  | Litter size | Mother ID |
|  |  |  |  | Age at death (months) |  |
| Adult | Weight | 2092 | 3860 | Sex | Year of capture |
|  |  |  |  | Population size | Permanent environment |
|  |  |  |  | Age (years) |  |
|  | Foreleg | 1936 | 3594 | Sex | Year of capture |
|  |  |  |  | Population size | Permanent environment |
|  |  |  |  | Age (years) |  |
|  | Hindleg | 2027 | 3481 | Sex | Year of capture |
|  |  |  |  | Population size | Permanent environment |
|  |  |  |  | Age (years) |  |
|  | Metacarpal | 987 | 987 | Sex | Year of birth |
|  |  |  |  | Age at death (years) |  |
|  | Jaw | 1057 | 1057 | Sex | Year of birth |
|  |  |  |  | Age at death (years) |  |

### Genetic data

8557 sheep have been genotyped on the Ovine SNP50 Illumina BeadChip, of which 38,130 SNPs are autosomal and polymorphic in the population. 188 individuals have additionally been genotyped on the Ovine Infinium HD SNP BeadChip which genotypes 600K SNPs – this allowed for imputation of the remaining genotyped individuals to this higher density. AlphaImpute v1.98 (Hickey et al. 2012) was used for the imputation as it combines shared haplotype and pedigree information to increase imputation accuracy (see Stoffel et al. 2021 for details on our imputation). Genotypes with a probability of < 0.99 were excluded, resulting in 419,281 autosomal SNPs remaining for 8557 individuals (4035 females, 4452 males). Imputed genotype “hard” calls were used instead of genotype probabilities in the analyses detailed in this manuscript. Locus positions for both sets of genetic data were based on the OAR_v3.1 genome assembly.

### Regional heritability mapping

Prior to regional heritability mapping, the traits were pre-corrected to account for genome-wide genetic diversity by fitting a LOCO GRM, constructed from all autosomes with the exception of the chromosome containing the focal region. We also fitted fixed and non-genetic random effects during pre-correction (see Table 1 for a full list of fixed and non-genetic random effects fitted). Pre-correction for the non-repeated measures traits was performed in DISSECT (Canela-Xandri et al. 2015) using the following model:

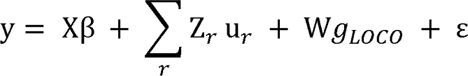

where y is the vector of phenotypic values; X is a design matrix linking individual records with the vector of fixed effects β, Z_r_ is an incidence matrix that relates a random effect to the individual records; u_r_ is the associated vector of non-genetic random effects; g_LOCO_ is the vector of additive genetic random effects from all autosomes except for that containing the focal region with W the incidence matrix linking individual phenotypes with the genetic effect; and ε is the vector of residuals. It is assumed that g_LOCO_ ∼*MVN*(0, Mσ_gLOCO_^2^) where σ_gLOCO_^2^ is the additive genetic variance explained by all autosomes except the excluded one, and M is the LOCO GRM.

The GRMs (VanRaden 2008) were computed using DISSECT, and the genetic relationship between individuals i and j is computed as:

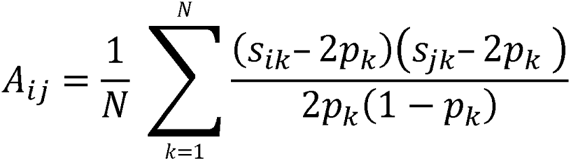

where s_ik_ is the number of copies of the reference allele for SNP k of the individual i, p_k_ is the frequency of the reference allele for the SNP k, and N is the number of SNPs.

The residual for each individual was then taken as the phenotype for RHM. Pre-correction for the three repeated measures traits (adult August weight, adult foreleg length and adult hindleg length) was performed using ASReml-R (version 4.1, Butler et al. 2017) using the same model as given above, and the mean of the residuals summed with the permanent environment effect for each individual was taken as the phenotype for RHM.

For all three regional heritability methods, we used the same regions to allow for direct comparisons between the methods. Due to Hap-RHM and SNHap-RHM requiring regions to be defined as haplotype blocks, we used haplotype blocks for all three methods. Haplotype blocks were estimated with Plink v1.90 using the *--blocks*, *--blocks-max-kb 500* (which allows pairs of variants within 500kb of each other to be considered within the same block) and *--blocks-min-maf 0.01* options (which instructs Plink to include all SNPs with a MAF higher than 0.01 when estimating the haplotype blocks (Purcell et al. 2007; Purcell 2014)). Using a higher max kb threshold or lower MAF threshold did not alter the haplotype block boundaries estimated. No haplotype block was allowed to have only one SNP, due to the SNP-based GRM and haplotype-based GRM being identical for such blocks. Any block containing only one SNP was therefore omitted from the analysis. Blocks were determined using all 8557 individuals with imputed genotypes to ensure consistency across phenotypes.

Phased data is required for Hap-RHM and SNHap-RHM; genotypes were phased using SHAPEIT v4.2 (Delaneau et al. 2019).

Regional heritability mapping was performed using the following models:

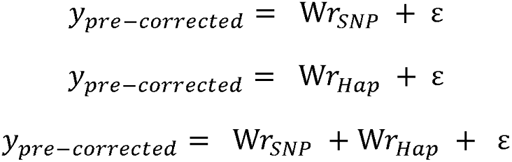

for SNP-RHM, Hap-RHM and SNHap-RHM respectively, where y_pre-corrected_ is the vector of pre-corrected phenotypic values, r_SNP_ is the vector of individual additive genetic random effects from all SNPs contained within the focal haplotype block and r_Hap_ is the vector of individual additive genetic random effects from the haplotype alleles for the focal haplotype block. It is assumed that r_SNP_ ∼ *MVN*(0, Mσ_rSNP_^2^) and r_Hap_ ∼ *MVN*(0, Mσ_rHap_^2^), where σ_rSNP_^2^ is the additive genetic variance from all SNPs in the haplotype block, σ_rHap_^2^ is the additive genetic variance from the haplotype alleles and M is the respective GRM. The GRMs were computed using DISSECT (Canela-Xandri et al. 2015). The SNP-based GRMs were calculated using the same method as the LOCO GRMs, except they were constructed from the SNPs located in the focal haplotype block. For the haplotype-based GRMs, the genetic relationship for individuals i and j is calculated as follows

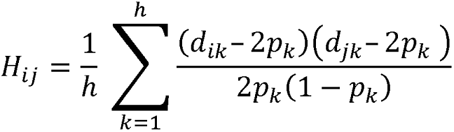

where *d_ik_* is the diplotype code (coded as the number of copies of haplotype *k*) for individual *i* and takes the values 0, 1, and 2, *pk* is the frequency of haplotype *k* and *h* is the number of haplotypes in the region (see Oppong et al. 2021 for further information and examples).

To test whether the regional heritability models explained significant variation for each region, we compared them against the null model:

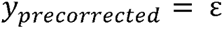

using loglikelihood ratio testing (LRT). We performed five comparisons; SNP-RHM, Hap-RHM and SNHap-RHM were all compared with the null model, and SNHap-RHM was additionally compared to each of SNP-RHM and Hap-RHM individually. LRTs were performed with 1 degree of freedom, with the exception of the comparison of SNHap-RHM to the null model, which was performed with 2 degrees of freedom. P values were calculated as 0.5× the p-value of a chi-squared distribution with one degree of freedom for the 1 degree of freedom tests. For the 2 degrees of freedom tests, the p values were calculated as 0.25× the p-value of a chi-squared distribution with two degrees of freedom plus 0.5× the p-value of a chi-squared distribution with one degree of freedom (Self and Liang 1987). Model fit was considered to be significantly improved if the resulting p value was less than 1.04e^-06^ (0.05 divided by 48,125, the total number of haplotype blocks).

### Comparison with GWAS

To determine how well the different RHM methods detected previously discovered loci, we identified which haplotype blocks contained the top SNP from each peak significantly associated with phenotypic variation for each trait when performing GWAS. GWAS and conditional GWAS analysis has recently been performed using the high density genotype data (James et al. 2022), so we used the results from that analysis. The significance threshold used in the GWAS analysis was 1.03e^−06^ (0.05/48635), which accounted for multiple testing using the SimpleM method (Gao et al. 2008). This method accounts for linkage disequilibrium between markers in order to calculate the effective number of independent tests.

### Identification of candidate genes

We extracted a list of genes overlapping any haplotype block for which model fit was improved by at least one RHM model, using the R biomaRt package (Durinck et al. 2005; Durinck et al. 2009) from the OAR_v3.1 genome assembly. Each gene was then reviewed against the Ensembl (Howe et al. 2020) and NCBI Gene (Bethesda (MD): National Library of Medicine (US) 2004 - 2023) databases to examine expression and functional annotations. Human and mouse orthologues were also used to characterise gene function due to the high level of genetic annotation in these two species.

## Results

### Soay sheep haplotype blocks

Setting the maximum kb between any two variants within the same haplotype block to 500Kb and the minimum minor allele frequency (MAF) for variants to be considered to 0.01 resulted in 48,125 haplotype blocks being estimated across the 26 Soay sheep autosomes. The maximum number of SNPs in a given haplotype block was 111, the minimum was 2 (as blocks with one SNP were omitted), and the average number of SNPs per haplotype was 8.19. 75% of haplotype blocks contained 10 or less SNPs, and 99% of blocks contained 50 or less. Block statistics for each chromosome are shown in Table 2.

**Table 2.** Number of SNPs per chromosome, number of haplotype blocks per chromosome, and average number of SNPs per haplotype block.

| Chromosome | Number of SNPs | Number of haplotype blocks | Mean number of SNPs per haplotype block |
| --- | --- | --- | --- |
| 1 | 47318 | 5466 | 8.66 |
| 2 | 41917 | 4426 | 9.47 |
| 3 | 38265 | 4081 | 9.38 |
| 4 | 20659 | 2259 | 9.15 |
| 5 | 18545 | 2132 | 8.70 |
| 6 | 19205 | 1908 | 10.07 |
| 7 | 17647 | 2267 | 7.78 |
| 8 | 15815 | 2029 | 7.79 |
| 9 | 16517 | 2177 | 7.59 |
| 10 | 15517 | 2223 | 6.98 |
| 11 | 11567 | 1371 | 8.44 |
| 12 | 13909 | 1665 | 8.35 |
| 13 | 13626 | 1427 | 9.55 |
| 14 | 10785 | 1205 | 8.95 |
| 15 | 13897 | 1537 | 9.04 |
| 16 | 11992 | 1400 | 8.57 |
| 17 | 12598 | 1694 | 7.44 |
| 18 | 11718 | 1357 | 8.64 |
| 19 | 10230 | 1058 | 9.67 |
| 20 | 9074 | 1115 | 8.14 |
| 21 | 7879 | 813 | 9.69 |
| 22 | 9352 | 1246 | 7.51 |
| 23 | 10004 | 1062 | 9.42 |
| 24 | 6362 | 492 | 12.93 |
| 25 | 7530 | 842 | 8.94 |
| 26 | 7353 | 873 | 8.42 |

Across the whole sample of genotyped individuals, haplotype allele frequency ranged from 0.00005843 to 0.9895992, with 0.00005843 being the most commonly observed haplotype frequency (28.95% of haplotype alleles). A frequency of 0.00005843 equates to a haplotype allele being present on one chromosome in the entire sample. 75% of haplotype alleles had a frequency lower than 0.072 (present on less than 1232 chromosomes in the entire sample), and 90% had a frequency lower than 0.316 (present on less than 5408 chromosomes in the entire sample) (Figure 1).

**Figure 1:**
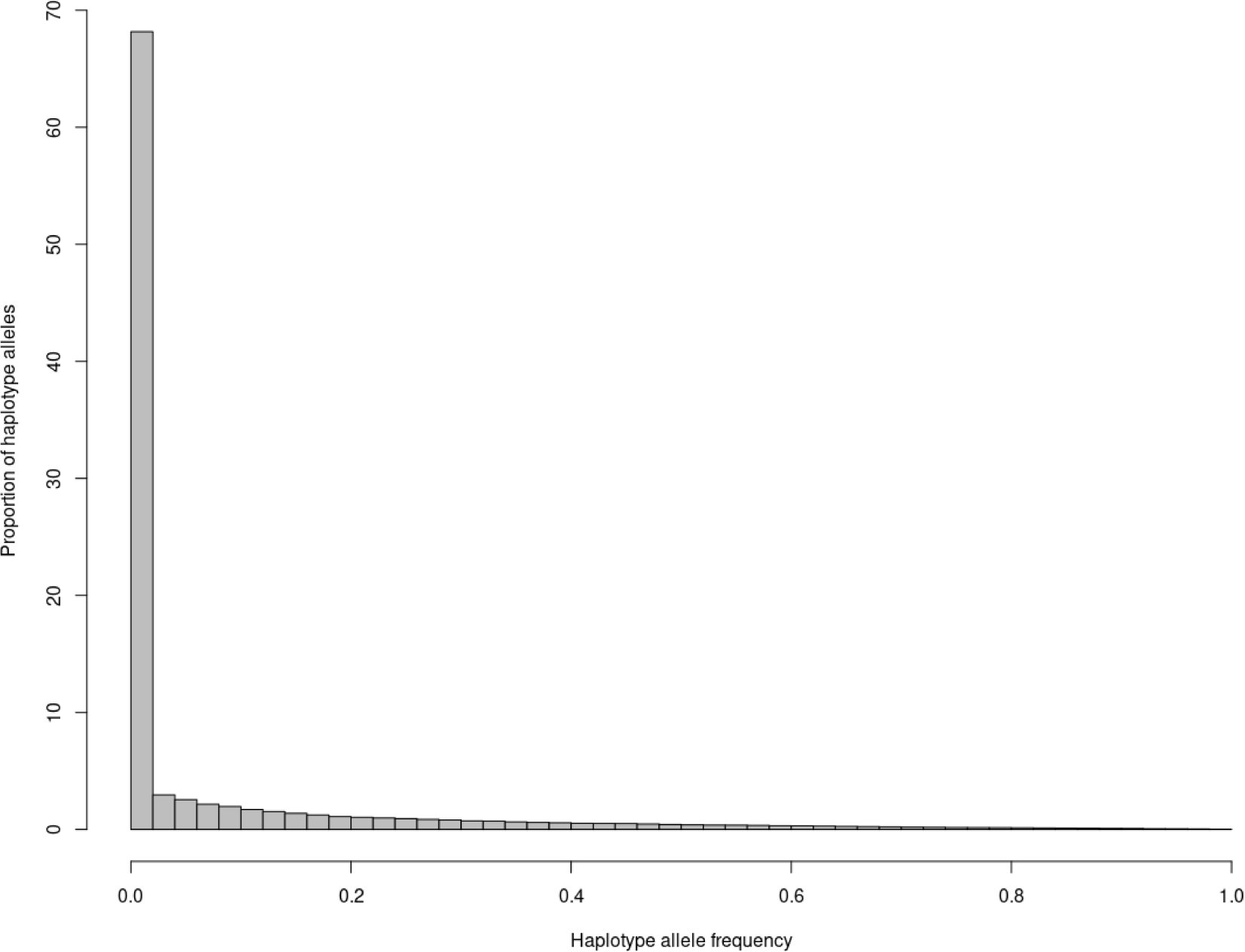
Proportion of haplotype alleles at each haplotype frequency over all regions.

### Comparison of RHM

A summary of results for the RHM analyses are shown in Table 3, whilst detailed results are shown in Supplementary Tables 1 – 10.

**Table 3.** Number of haplotype blocks for which inclusion of regional GRMs improved model fit. *SNP- RHM* column compares the SNP-RHM model against the null model to see if the inclusion of the regional SNP GRM improves model fit. *Hap-RHM* column compares the Hap-RHM model against the null model to see if the inclusion of the regional haplotype GRM improves model fit. *SNHap-RHM* column compares the SNHap-RHM model against the null model to see if the simultaneous inclusion of both the regional SNP GRM and the regional haplotype GRM improves model fit. *SNHap-RHM vs Hap-RHM* column compares the SNHap-RHM model against the Hap-RHM model to see if the additional inclusion of the regional SNP GRM improves model fit *S NHap-RHM vs SNP-RHM* column compares the SNHap-RHM model against the SNP-RHM model to see if the additional inclusion of the regional haplotype GRM improves model fit.

| Trait | SNP-RHM | Hap-RHM | SNHap-RHM | SNHap-RHM vs Hap-RHM | SNHap-RHM vs SNP-RHM |
| --- | --- | --- | --- | --- | --- |
| Birth weight | 0 | 0 | 0 | 0 | 0 |
| Lamb August weight | 0 | 0 | 0 | 0 | 0 |
| Lamb foreleg length | 0 | 2 | 0 | 0 | 0 |
| Lamb hindleg length | 0 | 2 | 0 | 0 | 0 |
| Lamb metacarpal length | 40 | 25 | 30 | 4 | 2 |
| Lamb jaw length | 0 | 5 | 0 | 0 | 2 |
| Adult August weight | 0 | 83 | 35 | 0 | 56 |
| Adult foreleg length | 0 | 6 | 6 | 0 | 6 |
| Adult hindleg length | 4 | 25 | 14 | 0 | 17 |
| Adult metacarpal length | 19 | 12 | 16 | 0 | 0 |
| Adult jaw length | 0 | 6 | 2 | 0 | 3 |

There were two traits for which none of the RHM models significantly improved model fit for any haplotype blocks: birth weight and lamb August weight, meaning that no regions of the genome were found to significantly explain additional genetic variance not accounted for during pre-correction.

For lamb foreleg length and lamb hindleg length, Hap-RHM was the only model which significantly improved model fit in comparison to the null model. Hap-RHM improved model fit for one haplotype block on chromosome 1 (haplotype number 1717) and one on chromosome 11 (726) for lamb foreleg length, and one on chromosome 2 (4160) and chromosome 3 (3432) for lamb hindleg length (Supplementary Tables 1 – 3).

For lamb metacarpal length, SNP-RHM significantly improved model fit for 16 haplotype blocks on chromosome 16. Model fit was improved for 14 of these blocks using Hap-RHM, with the other two being non-significant. 10 of the blocks for which model fit was improved by both SNP-RHM and Hap-RHM were also improved by SNHap-RHM when compared to the null model, however no blocks showed improved model fit when using SNHap-RHM when compared to either single-GRM model. For the same trait, SNP-RHM also improved model fit for 23 haplotype blocks on chromosome 19, whilst Hap-RHM improved model fit for 11 of the same blocks. When compared to the null model, SNHap-RHM improved model fit for 20 of these haplotype blocks on chromosome 19, and four haplotype blocks on chromosome 19 when compared to Hap-RHM. SNHap-RHM did not improve model fit for any blocks when compared to SNP-RHM (Supplementary Tables 1 and 4).

For lamb jaw length, model fit was only significantly improved by Hap-RHM and SNHap-RHM when compared to SNP-RHM. Hap-RHM improved model fit for one haplotype block on chromosome 3 (270), two blocks on chromosome 13 (1025 and 1041), one block on chromosome 14 (762) and one on chromosome 17 (923). When compared to SNP-RHM, SNHap-RHM significantly improved model fit for the same two blocks on chromosome 13 for which model fit was improved by Hap-RHM (Supplementary Tables 1 and 5).

For adult August weight, Hap-RHM significantly improved model fit for 83 haplotype blocks over 22 chromosomes. In comparison to the null model, SNHap-RHM improved model fit for 35 haplotype blocks over 21 chromosomes, whilst in comparison to SNP-RHM, model fit was improved for 56 haplotype blocks over 17 chromosomes. The blocks for which model fit was significantly improved by SNHap-RHM in comparison to either the null model or SNP-RHM were all ones that were also significantly improved by Hap-RHM, with the exception of one on chromosome 9 (1898), for which model fit was only significantly improved by SNHap-RHM in comparison to the null model. SNHap-RHM did not improve model fit for any haplotype blocks in comparison to Hap-RHM, nor did SNP- RHM when compared to the null model (Supplementary Tables 1 and 6).

For adult foreleg length, Hap-RHM significantly improved model fit for six haplotype blocks; one on chromosome 1 (1284), one on chromosome 6 (1064), one on chromosome 11 (113), one on chromosome 12 (23), one on chromosome 23 (1020) and one on chromosome 26 (259). Model fit was also improved for these same six haplotype blocks when comparing SNHap-RHM to the model and against SNP-RHM. SNHap-RHM did not improve model fit for any haplotype blocks in comparison to Hap-RHM, nor did SNP-RHM when compared to the null model (Supplementary Tables 1 and 7).

For adult hindleg length, SNP-RHM significantly improved model fit for four haplotype blocks on chromosome 16. Hap-RHM improved model fit for 25 haplotype blocks over 15 chromosomes, including two of the blocks for which model fit was significantly improved by SNP-RHM. SNHap-RHM improved model fit for 14 haplotype blocks over 11 chromosomes when compared to the null model, and 17 blocks over 12 chromosomes when compared to SNP-RHM. SNHap-RHM did not significantly improve model fit for any blocks when compared to Hap-RHM (Supplementary Tables 1 and 8).

For adult metacarpal length, SNP-RHM significantly improved model fit for 15 haplotype blocks on chromosome 16. Hap-RHM significantly improved model fit for 12 of these blocks. When compared to the null model, SNHap-RHM improved model fit for 12 haplotype blocks on chromosome 16 – all of which were blocks that experienced significant improvement in model fit by SNP-RHM. In addition, SNP-RHM and SNHap-RHM improved model fit for the same four blocks on chromosome 19 when compared to the null model. When compared to the single GRM RHM models, SNHap-RHM did not improve model fit for any haplotype blocks (Supplementary Tables 1 and 9).

For adult jaw length, Hap-RHM improved model fit for six haplotype blocks; one on chromosome 1 (3843), one on chromosome 3 (2046), one on chromosome 11 (1027), one on chromosome 18 (665) and two on chromosome 23 (342 and 434). Model fit was improved for the same haplotype blocks on chromosome 1 and 3 when comparing SNHap-RHM against both the null model and against SNP- RHM, with the haplotype block on chromosome 23 (434) also showing improved model fit when compared against SNP-RHM. SNHap-RHM did not improve model fit for any haplotype blocks in comparison to Hap-RHM, nor did SNP-RHM (Supplementary Tables 1 and 10).

### Comparison with GWAS results

Of the 11 traits, lamb August weight and lamb jaw length were the only two to have no previously associated genetic loci (Bérénos et al. 2014; James et al. 2022). Of the traits for which GWAS has previously identified SNP-trait associations, RHM only significantly improved model fit for blocks containing top SNPs associated with lamb metacarpal length, adult hindleg length, and adult metacarpal length on chromosomes 16 and 19.

The underlying causal variant on chromosome 16 influencing lamb metacarpal length is presumed to be the same variant influencing adult hindleg length and adult metacarpal length – the top GWAS SNP on chromosome 16 for adult hindleg length and lamb metacarpal length is the same (James et al. 2022), adult hindleg and metacarpal length have been shown to have a genetic correlation of 0.827 (S.E. 0.232) (Bérénos et al. 2014), and SNP-leg trait associations in this region have been shown to be dependent on each other; when a SNP genotype from this region is fitted during conditional analyses, no new SNP associations appear in this region. We can therefore combine the RHM results for these three traits to characterise the architecture of genetic variance in this region. Whilst SNP-RHM significantly improved model fit for blocks on chromosome 16 that Hap-RHM did not, there were no blocks on chromosome 16 for which Hap-RHM improved model fit but SNP-RHM did not (Supplementary Tables 1, 4, 8 and 9). In fact, in the case of adult hindleg length, Hap-RHM did not improve model fit for any blocks on chromosome 16 (Supplementary Table 8). This suggests that the additive genetic variance being attributed to the regional GRMs is due to individual SNP genotypes, rather than due to a specific haplotype allele. Block 1363 (which contains s22142.1, the top GWAS SNP for lamb metacarpal length and adult hindleg length) contains 17 SNPs and has 18 haplotype alleles in the population. The minor allele for s22142.1 appears in 3 haplotype alleles, with two of these haplotype alleles being relatively rare (each appearing on 17 chromosomes in the genotyped population).

Again, the underlying causal variant on chromosome 19 influencing lamb metacarpal length is presumed to be the same variant influencing adult metacarpal length – whilst the top GWAS SNPs are different for these two traits, they still fall in the same haplotype block (Supplementary Table 11), and when the genotype of the top SNP is fitted during conditional analysis, no new SNP-trait associations appear (James et al. 2022). Again, we can combine the RHM results for both lamb metacarpal length and adult metacarpal length to characterise the underlying architecture. For both traits, model fit for the block containing the top GWAS SNPs was only significantly improved by SNP- RHM and SNHap-RHM when compared to the null model (and SNHap-RHM compared to Hap-RHM in the case of lamb metacarpal length). This suggests that this association is being driven by the SNP alleles in this region, rather than the haplotype alleles. Block 952, which contains both top GWAS SNPs, has 37 SNPs and 52 haplotype alleles in the genotyped population. The minor alleles for each of the top SNPs each appear in two haplotype alleles, with one haplotype allele containing both minor SNP alleles. The haplotype alleles each containing one of the minor SNP alleles for the top GWAS SNPs were both rare in the population (appearing on one and 50 chromosomes in the population).

### Novel associations

Novel block-trait associations were identified using at least one RHM method for all but four traits – birth weight, lamb August weight, lamb metacarpal length and adult metacarpal length. In the case of the former two traits, RHM did not improve model fit for any haplotype blocks in comparison to the null model, whilst in the case of the latter two, RHM only significantly improved model fit in the same regions as previously identified QTL for these traits.

Across the lamb traits, at least one RHM method significantly improved model fit for a total of nine haplotype blocks; two blocks for lamb foreleg length (on chromosomes 1 and 11), two blocks for lamb foreleg length (on chromosomes 2 and 3), and five blocks for lamb jaw length (one on chromosome 3, two on chromosome 13, one on chromosome 14 and one on chromosome 17). For all of these trait-block associations, Hap-RHM was the only model to significantly improve model fit – with the exception of the two blocks on chromosome 13 for lamb jaw length, for which SNHap-RHM also improved model fit when compared to SNP-RHM.

At least one RHM method significantly improved model fit for adult August weight for 85 haplotype blocks that did not contain previously identified QTL across a total of 22 chromosomes. For the adult leg traits, at least one RHM method significantly improved model fit for multiple blocks that had not previously been identified via GWAS; six blocks across six chromosomes for adult foreleg length, and 23 blocks across 14 chromosomes for adult hindleg length respectively. For adult jaw length, model fit was significantly improved by at least one RHM method for six blocks across five chromosomes that had not previously been identified by GWAS. For all of these blocks, model fit was improved by a mixture of Hap-RHM, and SNHap-RHM when compared to either the null model or SNP-RHM.

### Genes in QTL regions

Across all haplotype blocks for which model fit was improved by at least one RHM method for at least one of the 11 focal traits, there were 351 genes overlapping these blocks. 91 of these genes are completely uncharacterised in sheep and classed as “novel genes”, and a further 14 genes were RNA genes. Of the 246 characterised protein coding genes, 13 genes had functional annotations that related to the traits for which model fit was improved (Table 4). One of these genes was in a haplotype block associated with lamb jaw length, four in haplotype blocks associated with adult August weight, one associated with adult foreleg and adult hindleg length, one in haplotype blocks associated only with adult hindleg length and eight in haplotype blocks associated with lamb metacarpal length (one of which was also associated with adult metacarpal length). One of these genes – *PTH1R* – was previously identified as a putative causal gene due to its functional data and proximity to top GWAS SNPs for multiple Soay sheep leg length measures (James et al. 2022).

**Table 4.**
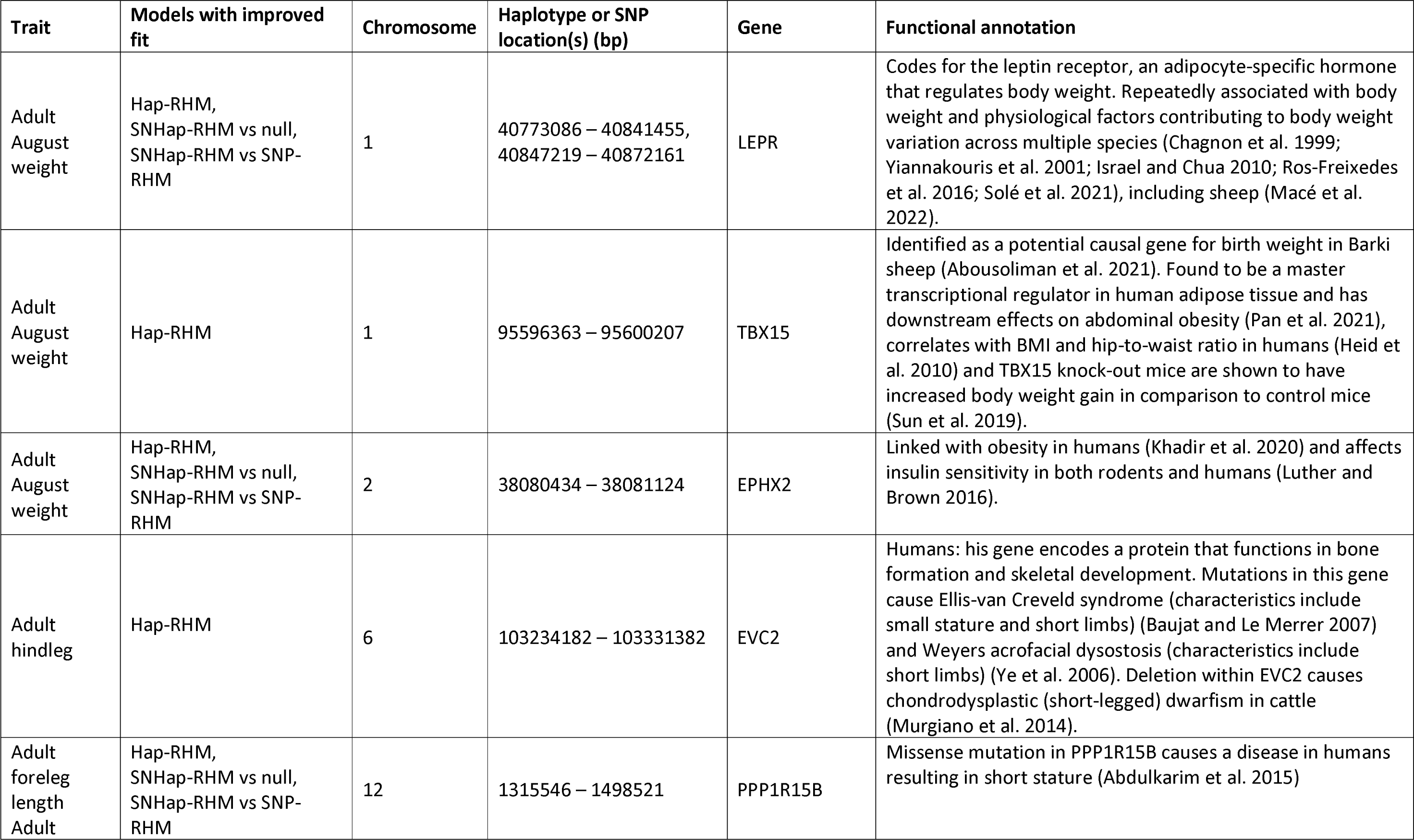
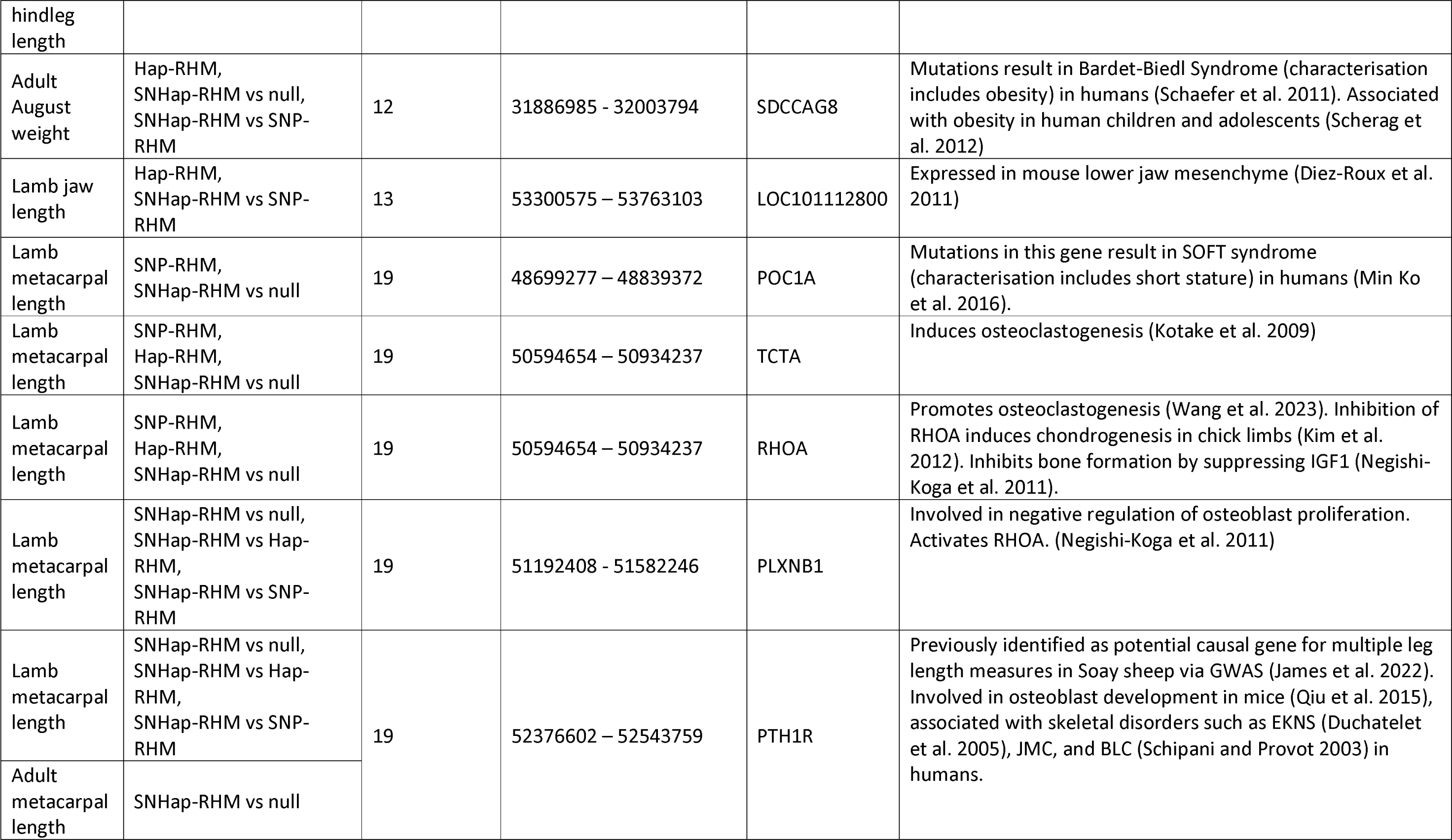
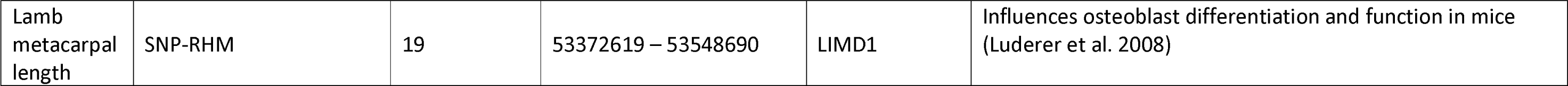
Potential candidate genes for future analyses. From left to right: associated trait, method that resulted in the gene being identified, chromosome, haplotype block, gene name, and evidence for association in sheep and other species.

## Discussion

### RHM overview

In total, there were 169 haplotype blocks for which model fit was improved for at least one trait by at least one RHM model. We found that Hap-RHM improved model fit more often than SNP-RHM, which is due in part to the fact that Hap-RHM improved model fit for more traits than SNP-RHM. Hap-RHM also improved model fit more often than SNHap-RHM when SNHap-RHM was compared to either the null model or either of the single regional GRM models. Additionally, SNP-RHM only improved model fit for haplotype blocks in regions surrounding QTL previously identified by GWAS.

We found that there were some trait associated regions identified via GWAS for which none of the RHM methods improved model fit – for instance, the regions on chromosomes 1 and 7 associated with birth weight, and the region on chromosome 16 associated with lamb foreleg length and hindleg length. There are two main differences between GWAS and RHM that may be contributing to the observed differences in results: firstly, SNP genotypes are fitted as fixed effects in GWAS, which confers more power than random effects. Secondly, we performed pre-correction of fixed and random effects prior to performing RHM but fitted them during the GWAS step.

We have previously shown that pre-correcting for fixed and random effects reduces power of GWAS to detect variant-trait associations. When pre-correction is performed for the adult traits, the only significant GWAS associations are those between SNPs on chromosome 16 and adult foreleg, hindleg and metacarpal lengths, and SNPs on chromosome 19 and adult metacarpal length (James et al. 2022). This mirrors our RHM results; the only haplotype blocks for which model fit was improved by the RHM methods were those two regions on chromosomes 16 and 19, with model fit for the latter region only improving for metacarpal length. Pre-correction may therefore explain why we did not see the RHM methods improving model fit for all of the haplotype blocks containing previously identified variants. Currently pre-correction is a necessary step when performing RHM with DISSECT due to DISSECT being unable to fit all of the necessary fixed and random effects during RHM. It would be interesting to rerun this analysis when suitable software is developed for single-step RHM, to determine whether single-step RHM improved model fit for all haplotype blocks containing significant GWAS associations.

The significance threshold used during GWAS in James et al. (2022) was similar to the threshold used during the RHM analyses (1.03e^-06^ and 1.04e^-06^ respectively). This is because we used SimpleM (Gao et al. 2008) to calculate the number of independent tests during GWAS – this was estimated to be 48,635, whilst the number of haplotype blocks estimated by Plink (Purcell et al. 2007; Purcell 2014) was 48,125. This is therefore not a major difference and won’t contribute to explaining why we obtained different results from GWAS and RHM.

RHM did, however, improve model fit for some regions associated with trait variation using GWAS; RHM improved model fit for haplotype blocks in regions previously found to be associated with lamb metacarpal length, adult hindleg length and adult metacarpal length on chromosome 16, and lamb metacarpal length and adult metacarpal length on chromosome 19.

SNP-RHM has previously been performed in a smaller sample of this same population, focusing on only adult morphometric traits (Bérénos et al. 2015). 37K autosomal SNPs were split into 150 SNP windows with a 75 SNP overlap. When comparing the results of (Bérénos et al. 2015) to our results for the same traits, we find six regions for which SNP-RHM improved model fit for (Bérénos et al. 2015) and at least one RHM method improved model fit in our own analyses; two regions on chromosome 1 and one region on chromosome 6 were associated with adult August weight (1:119,553,142 – 1:139,871,327, 1:163,370,112 – 1:173,759,083, 6:38,952,950 – 6:48762234), one region on chromosome 6 was associated with adult hindleg length (6:32,615,209 – 6:43,798,415), a region on chromsome 16 associated with adult hindleg and metacarpal length (16:64,064,879 – 16:71,555,691), and a region on chromosome 19 associated with adult metacarpal length (19:41,742,622 – 19:58,334,807).

### Genetic architecture of traits

As previously mentioned, RHM failed to significantly improve model fit for some haplotype blocks in regions previously associated with our focal traits by GWAS. However, RHM also identified some novel block-trait associations – for instance, at least one RHM method improved model fit for 85 haplotype blocks across a total of 22 chromosomes for adult August weight. None of these blocks were within 1Mb of a previously identified GWAS association. Interestingly, neither SNP-RHM compared to the null model nor SNHap-RHM when compared to Hap-RHM improved model fit for any haplotype blocks for adult August weight. This suggests that the majority of genetic variance contributing to variation in adult August weight is not due to small effect causal variants in LD with genotyped SNPs, but instead due to rare SNPs in LD with rare haplotype alleles or due to multiple SNPs in the same region interacting epistatically. We have previously shown that family-associated non-additive genetic variance such as dominance and epistasis may be making up 37.1% of previous narrow-sense heritability estimations for this trait (James et al. 2023). This finding would be consistent with Hap-RHM detecting regions in which multiple variants are acting in an epistasic manner. We found three genes with functional data suggesting an association with adult August weight that overlapped with haplotype blocks for which model fit was significantly improved by at least one RHM method: *LEPR*, *TBX15* and *EPHX2*. For the blocks overlapping all three genes, model fit is significantly improved by the presence of the haplotype GRM; Hap-RHM significantly improved model fit for all of the overlapping blocks, and SNHap-RHM significantly improved model fit when compared to the null model and to SNP-RHM for the blocks overlapping *LEPR* and *EPHX2*. This may explain why these regions were not identified as being associated with adult August weight when performing GWAS (James et al. 2022), as the variance influencing adult August weight in those regions is likely due to specific haplotype alleles, rather than individual SNP effects.

When performing RHM on lamb foreleg and hindleg lengths, only two blocks showed significantly improved model fit for each trait – none of which had previously been indicated to be associated with leg lengths in the Soay population. For all four blocks, Hap-RHM was the only model that significantly improved model fit, suggesting any variance being contributed by these blocks to their respective traits is due to the haplotypes, rather than individual SNPs within the blocks.

RHM improved model fit for a block on each of six chromosomes for adult foreleg length and 27 blocks across 15 chromosomes for adult hindleg length – of these only the blocks on chromosome 16 were close to top GWAS SNPs from any previous leg length GWAS results (Bérénos et al. 2014; James et al. 2022). Blocks on chromosome 16 overlapping top GWAS hits also showed improved model fit for both lamb and adult metacarpal length. It is hard to separate whether the variance being contributed by this region is due to individual SNP effects or an overall haplotype effect, as SNP- RHM, Hap-RHM and SNHap-RHM all significantly improved model fit when compared to the null model, but SNHap-RHM did not significantly improve model fit when compared to either of the single-GRM models. This does suggest, however, that the variance is not due to independent SNP and haplotype effects in the same block.

Blocks overlapping the top GWAS SNPs on chromosome 19 also showed significant improvement in model fit for lamb and adult metacarpal lengths when performing SNP-RHM and when comparing SNHap-RHM against the null model (and when compared to Hap-RHM for lamb metacarpal length). This suggests that the variance contributing to these traits is solely due to SNP genotypes, rather than haplotype alleles. Conditional analyses fitting the top GWAS SNP have shown that there are no secondary SNP-trait associations on chromosome 19 independent of the top SNP (James et al. 2022), implying that the variance is being contributed entirely by a single SNP.

RHM identified five block-trait associations for lamb jaw length. For all of these blocks, Hap-RHM was the only RHM model that improved model fit, with the exception of two blocks on chromosome 13 for which SNHap-RHM also improved model fit when compared to SNP-RHM. Similarly, six trait-block associations were identified for adult jaw length (none of which overlapped with lamb jaw length). Hap-RHM improved model fit for all six blocks, and SNHap-RHM improved model fit for two and three of these blocks when compared to the null model and SNP-RHM respectively. This suggests that these regions either contain a rare variant that influences jaw length that is only in LD with a low number of haplotype alleles, or they contain multiple variants interacting in an epistatic manner. It also implies that variation in lamb and adult jaw length are influenced by different genetic factors, which corroborates previous GWAS findings (James et al. 2022).

### Concluding remarks

We have demonstrated that RHM methods are a useful tool for detecting regions that contribute genetic variation to traits in a wild population and complement other analyses such as GWAS. We found that Hap-RHM and SNHap-RHM improved model fit for more haplotype blocks than SNP-RHM, but all three can be used together to better characterise the underlying genetic architecture within a region. Using these methods, we detected multiple haplotype blocks that improved model fit with at least one RHM method. From these regions, we characterised the genetic regions influencing trait variation and identified 13 potential genes that influence trait variation that have not previously been associated with variation in these traits in the Soay population.

