## Supplementary Table 1 for "Evaluating regional heritability mapping methods for identifying QTLs in a wild population of Soay sheep"

| Chromosome | Block | Block start (bp) | Block end (bp) | Block length (Kb) | SNPs | Haplotype alleles | Models | Genes | NCBI Gene name | Functional data |
|---|---|---|---|---|---|---|---|---|---|---|
| 1 | 948 | 40,773,086 | 40,841,455 | 68.37 | 10 | 14 | Adult August weight (Hap-RHM, SNHap-RHM vs null, SNHap-RHM vs SNP-RHM) | ENSOARG00000010394 | LEPR | Receptor for leptin (an adipocyte-specific hormone that regulates body weight). Mutations in this gene have been associated with obesity and pituitary dysfunction |
| 1 | 949 | 40,847,219 | 40,872,161 | 24.943 | 7 | 11 | Adult August weight (Hap-RHM) | ENSOARG00000010394 | LEPR | |
| 1 | 956 | 41,046,557 | 41,094,853 | 48.297 | 11 | 22 | Adult August weight (Hap-RHM, SNHap-RHM vs SNP-RHM) | | | |
| 1 | 974 | 41,690,586 | 41,727,381 | 36.796 | 10 | 28 | Adult August weight (Hap-RHM, SNHap-RHM vs null, SNHap-RHM vs SNP-RHM) | | | |
| 1 | 982 | 41,894,503 | 41,896,031 | 1.529 | 3 | 5 | Adult August weight (Hap-RHM, SNHap-RHM vs null, SNHap-RHM vs SNP-RHM) | | | |
| 1 | 1227 | 51,825,023 | 51,860,711 | 35.689 | 10 | 35 | Adult August weight (Hap-RHM) | | | |
| 1 | 1284 | 54,708,137 | 54,738,562 | 30.426 | 8 | 15 | Adult foreleg length (Hap-RHM, SNHap-RHM vs null, SNHap-RHM vs SNP-RHM) | ENSOARG00000013529 | ADGRL4 | |
| 1 | 1717 | 73,687,439 | 73,757,464 | 70.026 | 14 | 23 | Lamb foreleg length (Hap-RHM) | ENSOARG00000017549 | DPYD | |
| 1 | 1908 | 82,068,913 | 82,193,611 | 124.699 | 19 | 29 | Adult August weight (Hap-RHM, SNHap-RHM vs null, SNHap-RHM vs SNP-RHM) | | | |
| 1 | 2203 | 92,789,120 | 92,822,945 | 33.826 | 9 | 21 | Adult August weight (Hap-RHM, SNHap-RHM vs SNP-RHM) | | | |
| 1 | 2294 | 95,596,363 | 95,600,207 | 3.845 | 3 | 4 | Adult August weight (Hap-RHM) | ENSOARG00000020376 | TBX15 | Highly expressed in sheep muscle (Sheep tissue atlas). Mutations in humans associated with Cousin syndrome (a rare syndrome characterized mainly by short stature at birth, unusual facial appearance and skeletal abnormalities involving the shoulder blades and hips). High expression in fat in humans (HPA RNA-seq normal tissues). Adipose master transcription factor |
| 1 | 2542 | 106,366,046 | 106,386,067 | 20.022 | 3 | 10 | Adult August weight (Hap-RHM, SNHap-RHM vs null, SNHap-RHM vs SNP-RHM) | | | |
| 1 | 3115 | 139,277,588 | 139,295,840 | 18.253 | 8 | 17 | Adult August weight (Hap-RHM) | | | |
| 1 | 3587 | 160,031,383 | 160,095,549 | 64.167 | 7 | 8 | Adult August weight (Hap-RHM) | | | |
| 1 | 3749 | 171,350,897 | 171,455,010 | 104.114 | 16 | 21 | Adult August weight (Hap-RHM) | ENSOARG00000018897 | IFT57 | |
| 1 | 3843 | 178,080,084 | 178,183,944 | 103.861 | 15 | 27 | Adult jaw length (Hap-RHM, SNHap-RHM vs null, SNHap-RHM vs SNP-RHM) | | | |
| 1 | 3999 | 187,682,691 | 187,731,235 | 48.545 | 13 | 14 | Adult August weight (Hap-RHM, SNHap-RHM vs null, SNHap-RHM vs SNP-RHM) | ENSOARG00000020233 | HEG1 | |
| 1 | 5151 | 262,392,923 | 262,395,691 | 2.769 | 3 | 5 | Adult August weight (Hap-RHM) | ENSOARG00000011819 | AIRE | |
| 1 | 5152 | 262,407,194 | 262,416,861 | 9.668 | 6 | 11 | Adult August weight (Hap-RHM, SNHap-RHM vs null, SNHap-RHM vs SNP-RHM) | ENSOARG00000011864 | PFKL | |
| 1 | 5171 | 263,369,987 | 263,391,361 | 21.375 | 8 | 26 | Adult August weight (Hap-RHM, SNHap-RHM vs null, SNHap-RHM vs SNP-RHM) | ENSOARG00000012584 | COL18A1 | |
| 1 | 5173 | 263,421,836 | 263,469,212 | 47.377 | 10 | 31 | Adult August weight (Hap-RHM, SNHap-RHM vs SNP-RHM) | ENSOARG00000012584 | COL18A1 | |
| | | | | | | | | ENSOARG00000012653 | SLC19A1 | |
| 2 | 294 | 13,861,125 | 13,900,430 | 39.306 | 14 | 23 | Adult August weight (Hap-RHM, SNHap-RHM vs SNP-RHM) | ENSOARG00000006968 | PTPN3 | |
| 2 | 298 | 13,963,629 | 14,005,045 | 41.417 | 6 | 20 | Adult August weight (Hap-RHM, SNHap-RHM vs SNP-RHM) | ENSOARG00000007009 | EPB41L4B | |
| 2 | 847 | 38,080,434 | 38,081,124 | 0.691 | 2 | 4 | Adult August weight (Hap-RHM) | ENSOARG00000009659 | EPHX2 | Linked to obesity in humans |
| 2 | 1231 | 58,153,317 | 58,183,581 | 30.265 | 10 | 14 | Adult August weight (Hap-RHM, SNHap-RHM vs SNP-RHM) | ENSOARG00000012371 | CEP78 | |
| 2 | 1655 | 86,113,413 | 86,164,251 | 50.839 | 9 | 19 | Adult August weight (Hap-RHM) | ENSOARG00000021943 | U6 | |
| 2 | 1661 | 86,356,070 | 86,702,015 | 345.946 | 55 | 57 | Adult August weight (Hap-RHM, SNHap-RHM vs null, SNHap-RHM vs SNP-RHM) | ENSOARG00000014133 | ADAMTSL1 | |
| 2 | 1686 | 88,287,607 | 88,366,762 | 79.156 | 14 | 34 | Adult August weight (Hap-RHM, SNHap-RHM vs SNP-RHM) | ENSOARG00000014289 | | |
| 2 | 2219 | 136,485,236 | 136,485,296 | 0.061 | 2 | 4 | Adult August weight (Hap-RHM, SNHap-RHM vs null, SNHap-RHM vs SNP-RHM) | | | |
| 2 | 2876 | 164,795,365 | 165,011,337 | 215.973 | 31 | 27 | Adult August weight (Hap-RHM, SNHap-RHM vs SNP-RHM) | ENSOARG00000009898 | GTDC1 | |
| 2 | 4160 | 230,433,387 | 230,495,451 | 62.065 | 6 | 9 | Lamb hindleg length (Hap-RHM) | ENSOARG00000020608 | DNER | |
| | | | | | | | | ENSOARG00000022419 | U6 | |
| 3 | 270 | 15,517,976 | 15,665,753 | 147.778 | 30 | 39 | Lamb jaw length (Hap-RHM) | ENSOARG00000019363 | (novel gene) | |
| | | | | | | | | ENSOARG00000025958 | (novel gene) | |
| 3 | 366 | 20,227,521 | 20,376,157 | 148.637 | 36 | 45 | Adult August weight (Hap-RHM) | ENSOARG00000015997 | E2F6 | |
| | | | | | | | | ENSOARG00000022085 | 5s rRNA | |
| 3 | 1860 | 105,545,550 | 105,562,429 | 16.88 | 4 | 5 | Adult August weight (Hap-RHM) | | | |
| 3 | 2003 | 120,620,484 | 121,115,743 | 495.26 | 78 | 73 | Adult hindleg length (Hap-RHM, SNHap-RHM vs null, SNHap-RHM vs SNP-RHM) | ENSOARG00000015322 | SLC6A15 | |
| | | | | | | | | ENSOARG00000015341 | TSPAN19 | |
| | | | | | | | | ENSOARG00000023767 | U6 | |
| 3 | 2046 | 125,238,635 | 125,340,391 | 101.757 | 18 | 42 | Adult jaw length (Hap-RHM, SNHap-RHM vs null, SNHap-RHM vs SNP-RHM) | ENSOARG00000015607 | (novel gene) | |
| | | | | | | | | ENSOARG00000015617 | (novel gene) | |
| 3 | 2217 | 134,593,364 | 134,952,658 | 359.295 | 46 | 38 | Adult August weight (Hap-RHM, SNHap-RHM vs null, SNHap-RHM vs SNP-RHM) | ENSOARG00000017291 | SLC4A8 | |
| | | | | | | | | ENSOARG00000017314 | GALNT6 | |
| | | | | | | | | ENSOARG00000017327 | CELA1 | |
| | | | | | | | | ENSOARG00000017344 | BIN2 | |
| | | | | | | | | ENSOARG00000017355 | SMAGP | |
| | | | | | | | | ENSOARG00000017368 | DAZAP2 | |
| | | | | | | | | ENSOARG00000017381 | POU6F1 | |
| | | | | | | | | ENSOARG00000017390 | TFCP2 | |
| | | | | | | | | ENSOARG00000017411 | CSRNP2 | |
| | | | | | | | | ENSOARG00000017425 | LETMD1 | |
| 3 | 2749 | 163,548,241 | 163,642,434 | 94.194 | 22 | 24 | Adult hindleg length (Hap-RHM, SNHap-RHM vs SNP-RHM) | ENSOARG00000018012 | LOC101105903 | |
| | | | | | | | | ENSOARG00000018031 | LOC101107184 | |
| 3 | 2751 | 163,752,532 | 163,836,906 | 84.375 | 12 | 10 | Adult hindleg length (Hap-RHM, SNHap-RHM vs null, SNHap-RHM vs SNP-RHM) | ENSOARG00000011516 | (novel gene) | |
| 3 | 3432 | 199,253,201 | 199,277,862 | 24.662 | 8 | 14 | Lamb hindleg length (Hap-RHM) | ENSOARG00000020651 | PTPRO | |
| 3 | 4030 | 222,455,230 | 222,490,044 | 34.815 | 9 | 26 | Adult August weight (Hap-RHM, SNHap-RHM vs SNP-RHM) | | | |
| 3 | 4032 | 222,503,713 | 222,513,147 | 9.435 | 4 | 8 | Adult August weight (Hap-RHM) | | | |
| 4 | 268 | 13,003,091 | 13,088,097 | 85.007 | 12 | 24 | Adult August weight (Hap-RHM, SNHap-RHM vs SNP-RHM) | ENSOARG00000003766 | DYNC1I1 | |
| 4 | 345 | 16,637,915 | 16,645,702 | 7.788 | 3 | 6 | Adult August weight (Hap-RHM) | | | |
| 4 | 1770 | 97,010,982 | 97,103,764 | 92.783 | 21 | 32 | Adult hindleg length (Hap-RHM, SNHap-RHM vs null, SNHap-RHM vs SNP-RHM) | ENSOARG00000006769 | (novel gene) | |
| 6 | 213 | 12,129,893 | 12,195,170 | 65.278 | 12 | 21 | Adult August weight (Hap-RHM, SNHap-RHM vs null, SNHap-RHM vs SNP-RHM) | ENSOARG00000018416 | CAMK2D | |
| 6 | 405 | 19,545,275 | 19,571,212 | 25.938 | 6 | 11 | Adult August weight (Hap-RHM, SNHap-RHM vs null, SNHap-RHM vs SNP-RHM) | ENSOARG00000009956 | GSTCD | |

| | | | | | | | | | | |
|---|---|---|---|---|---|---|---|---|---|---|
| 6 | 676 | 40,594,243 | 40,598,722 | 4.48 | 4 | 9 | Adult August weight (Hap-RHM, SNHap-RHM vs null, SNHap-RHM vs SNP-RHM) | | | |
| 6 | 677 | 40,627,521 | 40,814,530 | 187.01 | 31 | 29 | Adult hindleg length (Hap-RHM, SNHap-RHM vs SNP-RHM) | | | |
| 6 | 810 | 52,709,410 | 52,878,546 | 169.137 | 23 | 40 | Adult August weight (Hap-RHM) | | | |
| 6 | 841 | 54,756,472 | 54,767,273 | 10.802 | 3 | 6 | Adult August weight (Hap-RHM, SNHap-RHM vs null, SNHap-RHM vs SNP-RHM) | | | |
| 6 | 855 | 55,360,902 | 55,466,024 | 105.123 | 16 | 41 | Adult August weight (Hap-RHM, SNHap-RHM vs SNP-RHM) | ENSOARG00000009322 | ARAP2 | |
| 6 | 1064 | 67,126,647 | 67,275,637 | 148.991 | 29 | 63 | Adult foreleg length (Hap-RHM, SNHap-RHM vs null, SNHap-RHM vs SNP-RHM) | ENSOARG00000017345 | SLC10A4 | |
| | | | | | | | | ENSOARG00000017393 | (novel gene) | |
| | | | | | | | | ENSOARG00000017797 | FRYL | |
| 6 | 1652 | 103,234,182 | 103,331,382 | 97.201 | 30 | 24 | Adult hindleg length (Hap-RHM) | ENSOARG00000007030 | EVC2 | Humans: his gene encodes a protein that functions in bone formation and skeletal development. Mutations in this gene, as well as in a neighboring gene that lies in a head-to-head configuration, cause Ellis-van Creveld syndrome |
| 7 | 1 | 24,558 | 209,889 | 185.332 | 19 | 142 | Adult August weight (Hap-RHM, SNHap-RHM vs null, SNHap-RHM vs SNP-RHM) | | | |
| 7 | 161 | 7,380,870 | 7,489,218 | 108.349 | 26 | 56 | Adult hindleg length (Hap-RHM, SNHap-RHM vs null, SNHap-RHM vs SNP-RHM) | ENSOARG00000016888 | IQGAP2 | |
| 7 | 179 | 8,344,719 | 8,375,455 | 30.737 | 6 | 22 | Adult August weight (Hap-RHM, SNHap-RHM vs null, SNHap-RHM vs SNP-RHM) | ENSOARG00000017022 | PDE8B | |
| | | | | | | | | ENSOARG00000026687 | (novel gene) | |
| 8 | 1689 | 78,683,108 | 78,734,898 | 51.791 | 9 | 9 | Adult August weight (Hap-RHM, SNHap-RHM vs SNP-RHM) | | | |
| 9 | 1381 | 57,985,117 | 58,057,584 | 72.468 | 12 | 16 | Adult August weight (Hap-RHM, SNHap-RHM vs SNP-RHM) | ENSOARG00000009590 | COLEC10 | |
| 9 | 1704 | 72,404,044 | 72,420,911 | 16.868 | 5 | 7 | Adult August weight (Hap-RHM) | ENSOARG00000026544 | (novel gene) | |
| | | | | | | | | ENSOARG00000026545 | (novel gene) | |
| 9 | 1723 | 73,438,286 | 73,446,206 | 7.921 | 4 | 7 | Adult August weight (Hap-RHM) | ENSOARG00000015979 | RIMS2 | |
| 9 | 1898 | 82,835,845 | 82,875,993 | 40.149 | 11 | 19 | Adult August weight (SNHap-RHM vs null) | | | |
| 9 | 2032 | 89,458,815 | 89,567,544 | 108.73 | 24 | 61 | Adult hindleg length (Hap-RHM, SNHap-RHM vs null, SNHap-RHM vs SNP-RHM) | | | |
| 10 | 483 | 18,080,744 | 18,096,127 | 15.384 | 5 | 11 | Adult August weight (Hap-RHM, SNHap-RHM vs null, SNHap-RHM vs SNP-RHM) | | | |
| 10 | 489 | 18,264,001 | 18,265,718 | 1.718 | 2 | 4 | Adult August weight (Hap-RHM, SNHap-RHM vs null, SNHap-RHM vs SNP-RHM) | | | |
| 10 | 491 | 18,306,871 | 18,342,967 | 36.097 | 7 | 16 | Adult August weight (Hap-RHM) | | | |
| 10 | 492 | 18,354,172 | 18,357,858 | 3.687 | 3 | 6 | Adult August weight (Hap-RHM) | | | |
| 10 | 790 | 29,520,015 | 29,535,013 | 14.999 | 3 | 6 | Adult August weight (Hap-RHM, SNHap-RHM vs SNP-RHM) | | | |
| 10 | 1194 | 50,858,554 | 50,876,668 | 18.115 | 7 | 20 | Adult August weight (Hap-RHM) | ENSOARG00000015582 | TBC1D4 | |
| 10 | 1587 | 63,549,495 | 63,585,886 | 36.392 | 6 | 9 | Adult August weight (Hap-RHM) | | | |
| 10 | 1628 | 65,020,171 | 65,157,788 | 137.618 | 14 | 14 | Adult August weight (Hap-RHM) | | | |
| 10 | 1629 | 65,167,961 | 65,227,127 | 59.167 | 10 | 16 | Adult August weight (Hap-RHM, SNHap-RHM vs SNP-RHM) | | | |
| 11 | 113 | 7,422,889 | 7,457,516 | 34.628 | 8 | 36 | Adult foreleg length (Hap-RHM, SNHap-RHM vs null, SNHap-RHM vs SNP-RHM), Adult hindleg length (Hap-RHM, SNHap-RHM vs null, SNHap-RHM vs SNP-RHM) | | | |
| 11 | 170 | 10,398,620 | 10,447,503 | 48.884 | 6 | 9 | Adult August weight (Hap-RHM, SNHap-RHM vs null, SNHap-RHM vs SNP-RHM) | ENSOARG00000014401 | RPS6KB1 | |
| | | | | | | | | ENSOARG00000014672 | RNFT1 | |
| | | | | | | | | ENSOARG00000023319 | (novel gene) | |
| 11 | 726 | 32,805,104 | 32,988,652 | 183.549 | 32 | 30 | Lamb foreleg length (Hap-RHM) | ENSOARG00000004399 | UBB | |
| | | | | | | | | ENSOARG00000017009 | (novel gene) | |
| | | | | | | | | ENSOARG00000017046 | PIGL | |
| | | | | | | | | ENSOARG00000017147 | NCOR1 | |
| 11 | 1027 | 50,247,507 | 50,268,735 | 21.229 | 7 | 9 | Adult jaw length (Hap-RHM) | | | |
| 12 | 23 | 1,315,546 | 1,498,521 | 182.976 | 23 | 106 | Adult foreleg length (Hap-RHM, SNHap-RHM vs null, SNHap-RHM vs SNP-RHM), Adult hindleg length (Hap-RHM, SNHap-RHM vs null, SNHap-RHM vs SNP-RHM) | ENSOARG00000002524 | PPP1R15B | PPP1R15B: Human: A Missense Mutation in PPP1R15B Causes a Syndrome Including Diabetes, Short Stature, and Microcephaly |
| | | | | | | | | ENSOARG00000002556 | (novel gene) | |
| | | | | | | | | ENSOARG00000002639 | PIK3C2B | |
| 12 | 696 | 31,886,985 | 32,003,794 | 116.81 | 19 | 23 | Adult August weight (Hap-RHM, SNHap-RHM vs SNP-RHM) | ENSOARG00000006316 | SDCCAG8 | Sdccag8: humans: linked to obesity |
| | | | | | | | | ENSOARG00000021882 | (novel gene) | |
| 12 | 737 | 35,365,159 | 35,408,696 | 43.538 | 13 | 70 | Adult August weight (Hap-RHM, SNHap-RHM vs null, SNHap-RHM vs SNP-RHM) | ENSOARG00000010135 | SLC19A2 | |
| | | | | | | | | ENSOARG00000010726 | SELP | |
| 12 | 1399 | 68,397,546 | 68,481,603 | 84.058 | 18 | 9 | Adult hindleg length (Hap-RHM) | ENSOARG00000009437 | RPS6KC1 | |
| 13 | 208 | 8,224,592 | 8,239,443 | 14.852 | 2 | 4 | Adult August weight (Hap-RHM, SNHap-RHM vs null, SNHap-RHM vs SNP-RHM) | | | |
| 13 | 212 | 8,321,481 | 8,340,545 | 19.065 | 4 | 6 | Adult August weight (Hap-RHM, SNHap-RHM vs null, SNHap-RHM vs SNP-RHM) | | | |
| 13 | 350 | 14,324,927 | 14,540,728 | 215.802 | 34 | 46 | Adult August weight (Hap-RHM, SNHap-RHM vs null, SNHap-RHM vs SNP-RHM) | | | |
| 13 | 488 | 21,753,962 | 21,782,926 | 28.965 | 4 | 6 | Adult August weight (Hap-RHM, SNHap-RHM vs SNP-RHM) | ENSOARG00000000512 | (novel gene) | |
| 13 | 1025 | 52,585,011 | 52,594,105 | 9.095 | 3 | 6 | Lamb jaw length (Hap-RHM, SNHap-RHM vs SNP-RHM) | | | |

| | | | | | | | | | | |
|---|---|---|---|---|---|---|---|---|---|---|
| 13 | 1041 | 53,300,575 | 53,763,103 | 462.529 | 54 | 49 | Lamb jaw length (Hap-RHM, SNHap-RHM vs SNP-RHM) | ENSOARG00000009948 | ZBTB46 | LOC101112800/Arf1gap: Mouse:Located in postsynaptic density. Is expressed in axial skeleton; craniocervical region bone; inner ear; and lower jaw mesenchyme |
| | | | | | | | | ENSOARG00000010128 | ZGPAT | |
| | | | | | | | | ENSOARG00000010188 | ARFRP1 | |
| | | | | | | | | ENSOARG00000010322 | TNFRSF6B | |
| | | | | | | | | ENSOARG00000010368 | RTEL1 | |
| | | | | | | | | ENSOARG00000010555 | STMN3 | |
| | | | | | | | | ENSOARG00000010596 | GMEB2 | |
| | | | | | | | | ENSOARG00000010686 | FNDC11 | |
| | | | | | | | | ENSOARG00000010780 | SRMS | |
| | | | | | | | | ENSOARG00000010950 | PTK6 | |
| | | | | | | | | ENSOARG00000010995 | (novel gene) | |
| | | | | | | | | ENSOARG00000011052 | EEF1A2 | |
| | | | | | | | | ENSOARG00000011107 | KCNQ2 | |
| | | | | | | | | ENSOARG00000011195 | CHRNA4 | |
| | | | | | | | | ENSOARG00000011265 | (novel gene) | |
| | | | | | | | | ENSOARG00000011406 | LOC101112800 | |
| | | | | | | | | ENSOARG00000011497 | (novel gene) | |
| | | | | | | | | ENSOARG00000011555 | (novel gene) | |
| | | | | | | | | ENSOARG00000011588 | BIRC7 | |
| | | | | | | | | ENSOARG00000011639 | YTHDF1 | |
| | | | | | | | | ENSOARG00000011782 | SLC17A9 | |
| | | | | | | | | ENSOARG00000022609 | (novel gene) | |
| | | | | | | | | ENSOARG00000026240 | (novel gene) | |
| 14 | 666 | 29,320,129 | 29,404,807 | 84.679 | 13 | 56 | Adult August weight (Hap-RHM, SNHap-RHM vs null, SNHap-RHM vs SNP-RHM) | | | |
| 14 | 762 | 34,218,884 | 34,718,824 | 499.941 | 59 | 19 | Lamb jaw length (Hap-RHM) | ENSOARG00000002860 | FHOD1 | |
| | | | | | | | | ENSOARG00000002877 | SLC9A5 | |
| | | | | | | | | ENSOARG00000002900 | PLEKHG4 | |
| | | | | | | | | ENSOARG00000002928 | KCTD19 | |
| | | | | | | | | ENSOARG00000002953 | LRRC36 | |
| | | | | | | | | ENSOARG00000002965 | TPPP3 | |
| | | | | | | | | ENSOARG00000002972 | ZDHHC1 | |
| | | | | | | | | ENSOARG00000002986 | HSD11B2 | |
| | | | | | | | | ENSOARG00000002996 | ATP6V0D1 | |
| | | | | | | | | ENSOARG00000003006 | AGRP | |
| | | | | | | | | ENSOARG00000003027 | RIPOR1 | |
| | | | | | | | | ENSOARG00000003053 | CTCF | |
| | | | | | | | | ENSOARG00000003079 | CARMIL2 | |
| | | | | | | | | ENSOARG00000003098 | ACD | |
| | | | | | | | | ENSOARG00000003109 | PARD6A | |
| | | | | | | | | ENSOARG00000003118 | ENKD1 | |
| | | | | | | | | ENSOARG00000003128 | C14H16orf86 | |
| | | | | | | | | ENSOARG00000003135 | GFOD2 | |
| | | | | | | | | ENSOARG00000003145 | RANBP10 | |
| | | | | | | | | ENSOARG00000003171 | TSNAXIP1 | |
| | | | | | | | | ENSOARG00000003186 | CENPT | |
| | | | | | | | | ENSOARG00000003196 | (novel gene) | |
| | | | | | | | | ENSOARG00000003200 | NUTF2 | |
| | | | | | | | | ENSOARG00000003220 | EDC4 | |
| 15 | 256 | 16,293,816 | 16,793,796 | 499.981 | 44 | 37 | Adult hindleg length (Hap-RHM, SNHap-RHM vs null, SNHap-RHM vs SNP-RHM) | ENSOARG00000010464 | ALKBH8 | |
| | | | | | | | | ENSOARG00000010653 | ELMOD1 | |
| | | | | | | | | ENSOARG00000016955 | SLN | |
| | | | | | | | | ENSOARG00000023352 | (novel gene) | |
| 15 | 1109 | 61,307,131 | 61,476,746 | 169.616 | 23 | 38 | Adult August weight (Hap-RHM, SNHap-RHM vs null, SNHap-RHM vs SNP-RHM) | ENSOARG00000016334 | WT1 | |
| | | | | | | | | ENSOARG00000021466 | (novel gene) | |
| | | | | | | | | ENSOARG00000021578 | (novel gene) | |
| | | | | | | | | ENSOARG00000022621 | (novel gene) | |
| | | | | | | | | ENSOARG00000023601 | (novel gene) | |
| | | | | | | | | ENSOARG00000024560 | (novel gene) | |
| | | | | | | | | ENSOARG00000024644 | (novel gene) | |
| 16 | 1326 | 68,447,430 | 68,454,153 | 6.724 | 3 | 4 | Lamb metacarpal length (SNP-RHM) | ENSOARG00000015559 | (novel gene) | |
| 16 | 1333 | 68,687,286 | 68,705,421 | 18.136 | 5 | 8 | Adult metacarpal length (SNP-RHM) | | | |
| 16 | 1340 | 68,915,832 | 68,981,502 | 65.671 | 13 | 27 | Lamb metacarpal length (SNP-RHM, Hap-RHM, SNHap-RHM vs null), Adult metacarpal length (SNP-RHM, Hap-RHM, SNHap-RHM vs null) | | | |
| 16 | 1346 | 69,129,929 | 69,138,953 | 9.025 | 6 | 12 | Lamb metacarpal length (SNP-RHM, Hap-RHM), Adult metacarpal length (SNP-RHM, Hap-RHM, SNHap-RHM vs null) | | | |
| 16 | 1347 | 69,151,199 | 69,158,314 | 7.116 | 3 | 5 | Lamb metacarpal length (SNP-RHM, Hap-RHM) | | | |
| 16 | 1348 | 69,160,398 | 69,181,657 | 21.26 | 6 | 11 | Lamb metacarpal length (SNP-RHM, Hap-RHM), Adult metacarpal length (SNP-RHM, Hap-RHM, SNHap-RHM vs null) | | | |
| 16 | 1352 | 69,226,093 | 69,306,236 | 80.144 | 18 | 29 | Lamb metacarpal length (SNP-RHM, Hap-RHM, SNHap-RHM vs null), Adult metacarpal length (SNP-RHM, Hap-RHM, SNHap-RHM vs null) | ENSOARG00000005781 | (novel gene) | |
| | | | | | | | | ENSOARG00000026985 | (novel gene) | |

| | | | | | | | | | | |
|---|---|---|---|---|---|---|---|---|---|---|
| 16 | 1360 | 69,425,086 | 69,574,670 | 149.585 | 30 | 32 | Lamb metacarpal length (SNP-RHM, Hap-RHM, SNHap-RHM vs null), | | | |
| | | | | | | | Adult metacarpal length (SNP-RHM, Hap-RHM, SNHap-RHM vs null) | | | |
| 16 | 1363 | 69,667,639 | 69,754,149 | 86.511 | 17 | 18 | Lamb metacarpal length (SNP-RHM, Hap-RHM, SNHap-RHM vs null), | | | |
| | | | | | | | Adult metacarpal length (SNP-RHM, Hap-RHM), | | | |
| | | | | | | | Adult hindleg length (SNP-RHM) | | | |
| 16 | 1367 | 69,839,345 | 69,853,299 | 13.955 | 6 | 9 | Lamb metacarpal length (SNP-RHM, Hap-RHM, SNHap-RHM vs null), | | | |
| | | | | | | | Adult metacarpal length (SNP-RHM, Hap-RHM, SNHap-RHM vs null), | | | |
| | | | | | | | Adult hindleg length (SNP-RHM, Hap-RHM, SNHap-RHM vs null) | | | |
| 16 | 1368 | 69,856,342 | 69,943,027 | 86.686 | 17 | 23 | Lamb metacarpal length (SNP-RHM, Hap-RHM), | | | |
| | | | | | | | Adult metacarpal length (SNP-RHM, Hap-RHM, SNHap-RHM vs null) | | | |
| 16 | 1379 | 70,307,056 | 70,368,449 | 61.394 | 15 | 17 | Lamb metacarpal length (SNP-RHM, Hap-RHM, SNHap-RHM vs null), | | | |
| | | | | | | | Adult metacarpal length (SNP-RHM, Hap-RHM, SNHap-RHM vs null) | | | |
| 16 | 1382 | 70,412,884 | 70,442,751 | 29.868 | 6 | 16 | Lamb metacarpal length (SNP-RHM, Hap-RHM, SNHap-RHM vs null), | | | |
| | | | | | | | Adult metacarpal length (SNP-RHM, Hap-RHM, SNHap-RHM vs null), | | | |
| | | | | | | | Adult hindleg length (SNP-RHM) | | | |
| 16 | 1384 | 70,469,116 | 70,480,771 | 11.656 | 5 | 6 | Adult metacarpal length (SNP-RHM, Hap-RHM, SNHap-RHM vs null) | ENSOARG00000015565 | (novel gene) | |
| | | | | | | | | ENSOARG00000015599 | CCDC127 | |
| | | | | | | | | ENSOARG00000026989 | (novel gene) | |
| 16 | 1385 | 70,492,788 | 70,861,290 | 368.503 | 32 | 62 | Lamb metacarpal length (SNP-RHM, Hap-RHM, SNHap-RHM vs null), | ENSOARG00000015619 | (novel gene) | |
| | | | | | | | Adult metacarpal length (SNP-RHM, SNHap-RHM vs null) | ENSOARG00000015678 | SDHA | |
| | | | | | | | | ENSOARG00000015729 | PDCD6 | |
| | | | | | | | | ENSOARG00000015756 | (novel gene) | |
| | | | | | | | | ENSOARG00000015791 | (novel gene) | |
| | | | | | | | | ENSOARG00000015862 | EXOC3 | |
| | | | | | | | | ENSOARG00000015903 | SLC9A3 | |
| | | | | | | | | ENSOARG00000015928 | CEP72 | |
| | | | | | | | | ENSOARG00000015975 | TPPP | |
| | | | | | | | | ENSOARG00000021220 | BRD9 | |
| | | | | | | | | ENSOARG00000026989 | 5S rRNA | |
| | | | | | | | | ENSOARG00000026990 | (novel gene) | |
| | | | | | | | | ENSOARG00000026991 | (novel gene) | |
| | | | | | | | | ENSOARG00000026992 | (novel gene) | |
| 16 | 1390 | 71,032,937 | 71,037,268 | 4.332 | 6 | 7 | Lamb metacarpal length (SNP-RHM, Hap-RHM, SNHap-RHM vs null), | ENSOARG00000016124 | (novel gene) | |
| | | | | | | | Adult hindleg length (SNP-RHM, Hap-RHM) | | | |
| 16 | 1393 | 71,117,303 | 71,134,260 | 16.958 | 4 | 9 | Lamb metacarpal length (SNP-RHM, Hap-RHM, SNHap-RHM vs null) | | | |
| 16 | 1394 | 71,135,107 | 71,417,145 | 282.039 | 24 | 39 | Lamb metacarpal length (SNP-RHM), | ENSOARG00000016135 | (novel gene) | |
| | | | | | | | Adult metacarpal length (SNP-RHM, SNHap-RHM vs null) | ENSOARG00000016170 | IRX4 | |
| | | | | | | | | ENSOARG00000021595 | 5S rRNA | |
| | | | | | | | | ENSOARG00000026994 | (novel gene) | |
| | | | | | | | | ENSOARG00000026995 | (novel gene) | |
| | | | | | | | | ENSOARG00000026996 | (novel gene) | |
| | | | | | | | | ENSOARG00000026997 | (novel gene) | |
| | | | | | | | | ENSOARG00000026998 | (novel gene) | |
| | | | | | | | | ENSOARG00000026999 | (novel gene) | |
| 16 | 1398 | 71,538,789 | 71,603,934 | 65.146 | 11 | 10 | Adult metacarpal length (SNP-RHM, Hap-RHM) | ENSOARG00000016211 | SLC6A3 | |
| | | | | | | | | ENSOARG00000027001 | (novel gene) | |
| 17 | 570 | 17,198,504 | 17,283,637 | 85.134 | 12 | 25 | Adult hindleg length (Hap-RHM) | ENSOARG00000012885 | (novel gene) | |
| 17 | 597 | 18,436,762 | 18,537,277 | 100.516 | 14 | 49 | Adult August weight (Hap-RHM, SNHap-RHM vs SNP-RHM) | ENSOARG00000025659 | (novel gene) | |
| 17 | 923 | 39,422,031 | 39,459,777 | 37.747 | 9 | 13 | Lamb jaw length (Hap-RHM) | ENSOARG00000002302 | RAPGEF2 | |
| 17 | 1368 | 57,691,208 | 57,747,777 | 56.57 | 11 | 15 | Adult hindleg length (Hap-RHM, SNHap-RHM vs null, SNHap-RHM vs SNP-RHM) | ENSOARG00000005170 | RNFT2 | |
| 17 | 1369 | 57,758,713 | 57,776,113 | 17.401 | 5 | 5 | Adult hindleg length (Hap-RHM) | ENSOARG00000004678 | (novel gene) | |
| 17 | 1669 | 71,033,562 | 71,045,469 | 11.908 | 2 | 4 | Adult August weight (Hap-RHM, SNHap-RHM vs null, SNHap-RHM vs SNP-RHM) | ENSOARG00000014555 | (novel gene) | |
| | | | | | | | | ENSOARG00000025708 | (novel gene) | |
| 17 | 1672 | 71,176,215 | 71,176,575 | 0.361 | 2 | 4 | Adult August weight (Hap-RHM) | ENSOARG00000015034 | (novel gene) | |
| 18 | 665 | 36,416,557 | 36,442,672 | 26.116 | 6 | 13 | Adult jaw length (Hap-RHM) | | | |
| 18 | 1248 | 62,650,233 | 62,714,362 | 64.13 | 14 | 41 | Adult hindleg length (Hap-RHM, SNHap-RHM vs null, SNHap-RHM vs SNP-RHM) | | | |
| 19 | 700 | 37,999,202 | 38,018,884 | 19.683 | 6 | 8 | Adult August weight (Hap-RHM, SNHap-RHM vs null, SNHap-RHM vs SNP-RHM) | ENSOARG00000011630 | SYNPR | |
| 19 | 915 | 48,504,989 | 48,559,115 | 54.127 | 12 | 19 | Lamb metacarpal length (SNP-RHM) | ENSOARG00000005161 | DNAH1 | |
| 19 | 921 | 48,699,277 | 48,839,372 | 140.096 | 19 | 22 | Lamb metacarpal length (SNP-RHM, SNHap-RHM vs null) | ENSOARG00000006352 | POC1A | POC1A: Mutations in this gene result in short stature, onychodysplasia, facial dysmorphism, and |
| | | | | | | | | ENSOARG00000006432 | DUSP7 | hypotrichosis (SOFT) syndrome. (Mouse) Acts upstream of or within several processes, including |
| | | | | | | | | ENSOARG00000026657 | (novel gene) | growth plate cartilage chondrocyte development |
| 19 | 923 | 48,882,534 | 48,959,233 | 76.7 | 22 | 32 | Lamb metacarpal length (SNP-RHM, SNHap-RHM vs null) | ENSOARG00000006456 | RPL29 | |
| | | | | | | | | ENSOARG00000006479 | ABHD14A | |
| | | | | | | | | ENSOARG00000006536 | ABHD14B | |
| | | | | | | | | ENSOARG00000006603 | PCBP4 | |
| | | | | | | | | ENSOARG00000006608 | (novel gene) | |
| | | | | | | | | ENSOARG00000006615 | (novel gene) | |
| | | | | | | | | ENSOARG00000006651 | (novel gene) | |
| | | | | | | | | ENSOARG00000006698 | RRP9 | |

| | | | | | | | | | | |
|---|---|---|---|---|---|---|---|---|---|---|
| 19 | 924 | 48,961,593 | 49,289,143 | 327.551 | 45 | 83 | Lamb metacarpal length (SNHap-RHM vs null, SNHap-RHM vs Hap-RHM) | ENSOARG00000006698 | RRP9 | |
| | | | | | | | | ENSOARG00000006825 | IQCF1 | |
| | | | | | | | | ENSOARG00000006874 | LOC101115831 | |
| | | | | | | | | ENSOARG00000006884 | IQCF2 | |
| | | | | | | | | ENSOARG00000006886 | IQCF3 | |
| | | | | | | | | ENSOARG00000006909 | (novel gene) | |
| | | | | | | | | ENSOARG00000006915 | IQCF6 | |
| | | | | | | | | ENSOARG00000006974 | GRM2 | |
| | | | | | | | | ENSOARG00000007079 | TEX264 | |
| | | | | | | | | ENSOARG00000007264 | RAD54L2 | |
| | | | | | | | | ENSOARG00000007546 | DCAF1 | |
| | | | | | | | | ENSOARG00000014150 | LOC101117539 | |
| | | | | | | | | ENSOARG00000023520 | U4 | |
| 19 | 926 | 49,334,367 | 49,391,499 | 57.133 | 7 | 10 | Lamb metacarpal length (SNP-RHM, Hap-RHM, SNHap-RHM vs null) | ENSOARG00000007757 | RBM15B | |
| | | | | | | | | ENSOARG00000007779 | MANF | |
| | | | | | | | | ENSOARG00000008062 | DOCK3 | |
| 19 | 931 | 49,807,150 | 49,823,282 | 16.133 | 6 | 10 | Lamb metacarpal length (SNP-RHM) | | | |
| 19 | 932 | 49,834,431 | 49,918,951 | 84.521 | 13 | 15 | Lamb metacarpal length (SNP-RHM, Hap-RHM, SNHap-RHM vs null) | ENSOARG00000009120 | CACNA2D2 | |
| 19 | 935 | 50,126,233 | 50,427,222 | 300.99 | 41 | 27 | Lamb metacarpal length (SNP-RHM, Hap-RHM, SNHap-RHM vs null) | ENSOARG00000010649 | SEMA3F | |
| | | | | | | | | ENSOARG00000010923 | RBM5 | |
| | | | | | | | | ENSOARG00000011125 | RBM6 | |
| | | | | | | | | ENSOARG00000011281 | MON1A | |
| | | | | | | | | ENSOARG00000011352 | MST1R | |
| | | | | | | | | ENSOARG00000011510 | CAMKV | |
| | | | | | | | | ENSOARG00000011570 | TRAIP | |
| | | | | | | | | ENSOARG00000011997 | UBA7 | |
| | | | | | | | | ENSOARG00000012365 | INKA1 | |
| | | | | | | | | ENSOARG00000012413 | IP6K1 | |
| | | | | | | | | ENSOARG00000012451 | GMPPB | |
| | | | | | | | | ENSOARG00000022525 | U6 | |
| 19 | 937 | 50,455,405 | 50,545,393 | 89.989 | 12 | 53 | Lamb metacarpal length (SNP-RHM, Hap-RHM, SNHap-RHM vs null) | ENSOARG00000013151 | MST1 | |
| | | | | | | | | ENSOARG00000013387 | APEH | |
| | | | | | | | | ENSOARG00000013564 | BSN | |
| 19 | 938 | 50,546,414 | 50,574,275 | 27.862 | 8 | 16 | Lamb metacarpal length (SNP-RHM, Hap-RHM, SNHap-RHM vs null) | ENSOARG00000013645 | DAG1 | |
| 19 | 940 | 50,594,654 | 50,934,237 | 339.584 | 50 | 25 | Lamb metacarpal length (SNP-RHM, Hap-RHM, SNHap-RHM vs null) | ENSOARG00000013728 | NICN1 | TCTA + RHOA : Humans + mice: Involved in negative regulation of osteoclast differentiation and |
| | | | | | | | | ENSOARG00000013773 | AMT | osteoclast fusion |
| | | | | | | | | ENSOARG00000013798 | TCTA | |
| | | | | | | | | ENSOARG00000013860 | RHOA | |
| | | | | | | | | ENSOARG00000013960 | USP4 | |
| | | | | | | | | ENSOARG00000014215 | C19H3orf62 | |
| | | | | | | | | ENSOARG00000014273 | IHO1 | |
| | | | | | | | | ENSOARG00000014296 | KLHDC8B | |
| | | | | | | | | ENSOARG00000014388 | CCDC71 | |
| | | | | | | | | ENSOARG00000014559 | LOC101106719 | |
| | | | | | | | | ENSOARG00000014927 | USP19 | |
| | | | | | | | | ENSOARG00000015195 | QARS1 | |
| | | | | | | | | ENSOARG00000015363 | QRICH1 | |
| | | | | | | | | ENSOARG00000015447 | IMPDH2 | |
| | | | | | | | | ENSOARG00000015657 | NDUFAF3 | |
| | | | | | | | | ENSOARG00000015672 | DALRD3 | |
| | | | | | | | | ENSOARG00000015805 | WDR6 | |
| | | | | | | | | ENSOARG00000015842 | P4HTM | |
| | | | | | | | | ENSOARG00000022437 | (novel gene) | |
| | | | | | | | | ENSOARG00000023735 | Oar-mir-191 | |
| | | | | | | | | ENSOARG00000023885 | U6 | |
| 19 | 941 | 50,982,456 | 51,125,248 | 142.793 | 23 | 30 | Lamb metacarpal length (SNP-RHM, SNHap-RHM vs null) | ENSOARG00000016084 | SLC25A20 | |
| | | | | | | | | ENSOARG00000016132 | PRKAR2A | |
| 19 | 942 | 51,143,353 | 51,144,906 | 1.554 | 4 | 4 | Lamb metacarpal length (SNP-RHM, Hap-RHM) | ENSOARG00000016244 | NCKIPSD | |
| 19 | 943 | 51,154,170 | 51,171,583 | 17.414 | 3 | 5 | Lamb metacarpal length (SNP-RHM, Hap-RHM, SNHap-RHM vs null) | ENSOARG00000016244 | NCKIPSD | |
| | | | | | | | | ENSOARG00000016424 | CELSR3 | |

| | | | | | | | | | | |
|---|---|---|---|---|---|---|---|---|---|---|
| 19 | 945 | 51,192,408 | 51,582,246 | 389.839 | 52 | 60 | Lamb metacarpal length (SNHap-RHM vs null, SNHap-RHM vs Hap-RHM) | ENSOARG00000000092 | CELSR3 | PLXNB1: Mouse: Involved in several processes, including negative regulation of osteoblast proliferation |
| | | | | | | | | ENSOARG00000000153 | SHISA5 | |
| | | | | | | | | ENSOARG00000000255 | TREX1 | |
| | | | | | | | | ENSOARG00000000409 | ATRIP | |
| | | | | | | | | ENSOARG00000000605 | CCDC51 | |
| | | | | | | | | ENSOARG00000000999 | PLXNB1 | |
| | | | | | | | | ENSOARG00000001093 | SPINK8 | |
| | | | | | | | | ENSOARG00000001285 | NME6 | |
| | | | | | | | | ENSOARG00000001392 | LOC101112427 | |
| | | | | | | | | ENSOARG00000001481 | (novel gene) | |
| | | | | | | | | ENSOARG00000001605 | (novel gene) | |
| | | | | | | | | ENSOARG00000001802 | LOC105607776 | |
| | | | | | | | | ENSOARG00000001969 | CATHL3 | |
| | | | | | | | | ENSOARG00000002027 | BAC5 | |
| | | | | | | | | ENSOARG00000014171 | (novel gene) | |
| | | | | | | | | ENSOARG00000016424 | UCN2 | |
| | | | | | | | | ENSOARG00000016539 | SLC26A6 | |
| | | | | | | | | ENSOARG00000016645 | TMEM89 | |
| | | | | | | | | ENSOARG00000016718 | LOC101110095 | |
| | | | | | | | | ENSOARG00000017257 | COL7A1 | |
| | | | | | | | | ENSOARG00000017897 | PFKFB4 | |
| | | | | | | | | ENSOARG00000021670 | (novel gene) | |
| | | | | | | | | ENSOARG00000026659 | (novel gene) | |
| | | | | | | | | ENSOARG00000026660 | (novel gene) | |
| | | | | | | | | ENSOARG00000026661 | (novel gene) | |
| 19 | 946 | 51,583,587 | 51,641,462 | 57.876 | 14 | 22 | Lamb metacarpal length (SNP-RHM, Hap-RHM, SNHap-RHM vs null) | ENSOARG00000002154 | SC5 | |
| | | | | | | | | ENSOARG00000002373 | CDC25A | |
| | | | | | | | | ENSOARG00000022608 | 5S rRNA | |
| 19 | 948 | 51,665,021 | 51,881,263 | 216.243 | 40 | 64 | Lamb metacarpal length (SNP-RHM, SNHap-RHM vs null), Adult metacarpal length (SNP-RHM, SNHap-RHM vs null) | ENSOARG00000002647 | MAP4 | |
| | | | | | | | | ENSOARG00000002814 | (novel gene) | |
| | | | | | | | | ENSOARG00000002897 | DHX30 | |
| | | | | | | | | ENSOARG00000024996 | (novel gene) | |
| 19 | 950 | 51,908,871 | 52,340,442 | 431.572 | 66 | 72 | Lamb metacarpal length (SNP-RHM, SNHap-RHM vs null, SNHap-RHM vs Hap-RHM), Adult metacarpal length (SNP-RHM, SNHap-RHM vs null) | ENSOARG00000003267 | SMARCC1 | |
| | | | | | | | | ENSOARG00000003512 | CSPG5 | |
| | | | | | | | | ENSOARG00000003560 | ELP6 | |
| | | | | | | | | ENSOARG00000003881 | SCAP | |
| | | | | | | | | ENSOARG00000004081 | (novel gene) | |
| | | | | | | | | ENSOARG00000004151 | LOC101122819 | |
| | | | | | | | | ENSOARG00000004308 | KLHL18 | |
| | | | | | | | | ENSOARG00000004598 | KIF9 | |
| | | | | | | | | ENSOARG00000024491 | U6 | |
| | | | | | | | | ENSOARG00000026662 | (novel gene) | |
| 19 | 952 | 52,376,602 | 52,543,759 | 167.158 | 37 | 52 | Lamb metacarpal length (SNP-RHM, SNHap-RHM vs null, SNHap-RHM vs Hap-RHM), Adult metacarpal length (SNP-RHM, SNHap-RHM vs null) | ENSOARG00000005054 | SETD2 | PTH1R: Humans: Defects in this receptor are known to be the cause of Jansen's metaphyseal chondrodysplasia (JMC), chondrodysplasia Blomstrand type (BOCD), as well as enchodromatosis. In bone, it is expressed on the surface of osteoblasts. When the receptor is activated through PTH binding, osteoblasts express RANKL (Receptor Activator of Nuclear Factor kB Ligand), which binds to RANK (Receptor Activator of Nuclear Factor kB) on osteoclasts. This turns on osteoclasts to ultimately increase the resorption rate. |
| | | | | | | | | ENSOARG00000005435 | LOC101114909 | |
| | | | | | | | | ENSOARG00000005857 | NBEAL2 | |
| | | | | | | | | ENSOARG00000006426 | CCDC12 | |
| | | | | | | | | ENSOARG00000006638 | PTH1R | |
| 19 | 954 | 52,570,585 | 52,595,896 | 25.312 | 7 | 10 | Lamb metacarpal length (SNP-RHM, SNHap-RHM vs null) | ENSOARG00000007025 | LOC101115659 | |
| 19 | 957 | 52,659,885 | 52,732,395 | 72.511 | 20 | 28 | Lamb metacarpal length (SNP-RHM, Hap-RHM, SNHap-RHM vs null) | ENSOARG00000007290 | LOC101102765 | |
| | | | | | | | | ENSOARG00000007381 | PRSS50 | |
| | | | | | | | | ENSOARG00000007470 | TMIE | |
| | | | | | | | | ENSOARG00000007704 | (novel gene) | |
| | | | | | | | | ENSOARG00000014187 | (novel gene) | |
| | | | | | | | | ENSOARG00000026663 | ALS2CL | |
| 19 | 958 | 52,733,279 | 52,798,829 | 65.551 | 15 | 20 | Lamb metacarpal length (SNP-RHM, Hap-RHM, SNHap-RHM vs null), Adult metacarpal length (SNP-RHM, SNHap-RHM vs null) | ENSOARG00000007704 | ALS2CL | |
| | | | | | | | | ENSOARG00000008087 | FAM240A | |
| | | | | | | | | ENSOARG00000008117 | CRIPTO | |
| 19 | 964 | 52,912,220 | 53,099,344 | 187.125 | 37 | 45 | Lamb metacarpal length (SNP-RHM, SNHap-RHM vs null) | ENSOARG00000014203 | CCRL2 | |
| | | | | | | | | ENSOARG00000014214 | CCR5 | |
| | | | | | | | | ENSOARG00000014236 | LOC101117706 | |
| | | | | | | | | ENSOARG00000014244 | CCR3 | |
| 19 | 969 | 53,372,619 | 53,548,690 | 176.072 | 29 | 52 | Lamb metacarpal length (SNP-RHM) | ENSOARG00000009571 | LZTFL1 | LIMD1: Mouse: Involved in negative regulation of canonical Wnt signaling pathway; negative regulation of osteoblast differentiation; and osteoblast development. Expressed in limb |
| | | | | | | | | ENSOARG00000009813 | SLC6A20 | |
| | | | | | | | | ENSOARG00000010162 | SACM1L | |
| | | | | | | | | ENSOARG00000010390 | LIMD1 | |
| 20 | 3 | 358,178 | 802,423 | 444.246 | 42 | 143 | Adult hindleg length (Hap-RHM, SNHap-RHM vs null, SNHap-RHM vs SNP-RHM) | ENSOARG00000005603 | KHDRBS2 | |
| | | | | | | | | ENSOARG00000024249 | U6 | |
| 20 | 8 | 1,254,840 | 1,315,189 | 60.35 | 13 | 13 | Adult hindleg length (Hap-RHM, SNHap-RHM vs null, SNHap-RHM vs SNP-RHM) | | | |
| 20 | 263 | 14,097,841 | 14,118,776 | 20.936 | 6 | 8 | Adult August weight (Hap-RHM, SNHap-RHM vs SNP-RHM) | | | |

| | | | | | | | | | |
|---|---|---|---|---|---|---|---|---|---|
| 20 | 273 | 14,428,550 | 14,485,702 | 57.153 | 10 | 16 | Adult August weight (Hap-RHM) | | |
| 20 | 1059 | 47,667,243 | 47,726,425 | 59.183 | 10 | 22 | Adult hindleg length (Hap-RHM) | | |
| 20 | 1075 | 48,121,607 | 48,242,296 | 120.69 | 24 | 63 | Adult hindleg length (Hap-RHM, SNHap-RHM vs SNP-RHM) | ENSOARG00000018718 | (novel gene) |
| 21 | 307 | 13,200,687 | 13,254,516 | 53.83 | 8 | 10 | Adult hindleg length (Hap-RHM) | | |
| 21 | 746 | 45,263,507 | 45,345,862 | 82.356 | 23 | 19 | Adult August weight (Hap-RHM) | ENSOARG00000015793 | PPP6R3 |
| 22 | 138 | 8,349,025 | 8,370,713 | 21.689 | 5 | 12 | Adult August weight (Hap-RHM, SNHap-RHM vs null, SNHap-RHM vs SNP-RHM) | | |
| 22 | 140 | 8,379,465 | 8,400,165 | 20.701 | 5 | 11 | Adult August weight (Hap-RHM, SNHap-RHM vs SNP-RHM) | ENSOARG00000013806 | ASAH2 |
| 22 | 327 | 13,291,916 | 13,300,911 | 8.996 | 3 | 6 | Adult August weight (Hap-RHM) | ENSOARG00000016671 | CPEB3 |
| 22 | 1167 | 44,452,770 | 44,548,081 | 95.312 | 21 | 52 | Adult August weight (Hap-RHM) | ENSOARG00000012241 | ADAM12 |
| 22 | 1246 | 50,486,318 | 50,820,764 | 334.447 | 43 | 62 | Adult August weight (Hap-RHM, SNHap-RHM vs SNP-RHM) | ENSOARG00000000042 | CYP2E1 |
| | | | | | | | | ENSOARG00000000369 | ECHS1 |
| | | | | | | | | ENSOARG00000000663 | FUOM |
| | | | | | | | | ENSOARG00000000727 | (novel gene) |
| | | | | | | | | ENSOARG00000000817 | CALY |
| | | | | | | | | ENSOARG00000000965 | ZNF511 |
| | | | | | | | | ENSOARG00000001302 | TUBGCP2 |
| | | | | | | | | ENSOARG00000014956 | (novel gene) |
| | | | | | | | | ENSOARG00000014969 | (novel gene) |
| | | | | | | | | ENSOARG00000014978 | (novel gene) |
| | | | | | | | | ENSOARG00000017206 | CFAP46 |
| | | | | | | | | ENSOARG00000018365 | (novel gene) |
| | | | | | | | | ENSOARG00000018459 | MTG1 |
| | | | | | | | | ENSOARG00000018583 | (novel gene) |
| | | | | | | | | ENSOARG00000018620 | (novel gene) |
| | | | | | | | | ENSOARG00000026777 | (novel gene) |
| | | | | | | | | ENSOARG00000026779 | (novel gene) |
| 23 | 342 | 14,087,536 | 14,348,846 | 261.311 | 13 | 7 | Adult jaw length (Hap-RHM) | ENSOARG00000003785 | (novel gene) |
| | | | | | | | | ENSOARG00000005184 | PIK3C3 |
| 23 | 434 | 21,076,868 | 21,139,467 | 62.6 | 12 | 32 | Adult jaw length (Hap-RHM, SNHap-RHM vs SNP-RHM) | ENSOARG00000021861 | U6 |
| 23 | 642 | 39,064,510 | 39,116,473 | 51.964 | 10 | 15 | Adult August weight (Hap-RHM, SNHap-RHM vs null, SNHap-RHM vs SNP-RHM) | ENSOARG00000026186 | (novel gene) |
| 23 | 643 | 39,126,867 | 39,137,482 | 10.616 | 4 | 5 | Adult August weight (Hap-RHM, SNHap-RHM vs null, SNHap-RHM vs SNP-RHM) | ENSOARG00000026186 | (novel gene) |
| 23 | 651 | 39,387,808 | 39,419,963 | 32.156 | 6 | 11 | Adult August weight (Hap-RHM) | | |
| 23 | 667 | 39,908,616 | 39,931,884 | 23.269 | 8 | 13 | Adult August weight (Hap-RHM, SNHap-RHM vs null, SNHap-RHM vs SNP-RHM) | | |
| 23 | 673 | 40,052,504 | 40,140,303 | 87.8 | 16 | 34 | Adult August weight (Hap-RHM, SNHap-RHM vs null, SNHap-RHM vs SNP-RHM) | ENSOARG00000000085 | ARHGAP28 |
| | | | | | | | | ENSOARG00000026188 | (novel gene) |
| 23 | 1020 | 58,109,141 | 58,167,269 | 58.129 | 12 | 61 | Adult foreleg length (Hap-RHM, SNHap-RHM vs null, SNHap-RHM vs SNP-RHM) | ENSOARG00000005569 | ZNF532 |
| 24 | 489 | 41,675,204 | 41,759,916 | 84.713 | 9 | 36 | Adult August weight (Hap-RHM, SNHap-RHM vs null, SNHap-RHM vs SNP-RHM) | ENSOARG00000006153 | PDGFA |
| | | | | | | | | ENSOARG00000025920 | (novel gene) |
| 25 | 82 | 4,586,431 | 4,883,217 | 296.787 | 50 | 50 | Adult August weight (Hap-RHM, SNHap-RHM vs SNP-RHM) | ENSOARG00000003291 | DISC1 |
| | | | | | | | | ENSOARG00000023491 | (novel gene) |
| 25 | 839 | 45,065,687 | 45,210,626 | 144.94 | 30 | 319 | Adult hindleg length (Hap-RHM, SNHap-RHM vs SNP-RHM) | ENSOARG00000002999 | ZNF32 |
| | | | | | | | | ENSOARG00000026418 | (novel gene) |
| 26 | 259 | 13,997,630 | 14,095,506 | 97.877 | 13 | 31 | Adult foreleg length (Hap-RHM, SNHap-RHM vs null, SNHap-RHM vs SNP-RHM) | | |
| 26 | 288 | 15,931,720 | 15,994,925 | 63.206 | 11 | 33 | Adult hindleg length (Hap-RHM, SNHap-RHM vs null, SNHap-RHM vs SNP-RHM) | | |

Supplementary Table 1 – Full list of haplotype blocks for which RHM significantly improved model fit for at least one trait. From left to right: chromosome, haplotype block number, haplotype block start (bp), haplotype block end (bp), length of haplotype block (Kb), number of SNPs in the block, number of haplotype alleles in the block, the trait-RHM model combinations that showed significant improvement in model fit, the Ensembl gene IDs of the genes located in or overlapping the block, the NCBI gene names, and any functional data relevant to the traits.
