## Supplementary Table 5 for "Evaluating regional heritability mapping methods for identifying QTLs in a wild population of Soay sheep"

| Chromosome | Block start (bp) | Block end (bp) | SNP-RHM | Hap-RHM | SNHap-RHM vs null | SNHap-RHM vs Hap-RHM | SNHap-RHM vs SNP-RHM |
|---|---|---|---|---|---|---|---|
| 3 | 15,517,976 | 15,665,753 | Not significant | 4.93E-07 | Model failed | Model failed | Model failed |
| 13 | 52,585,011 | 52,594,105 | Not significant | 2.49E-07 | Not significant | Not significant | 5.20E-07 |
| 13 | 53,300,575 | 53,763,103 | Not significant | 5.08E-07 | Not significant | Not significant | 5.46E-07 |
| 14 | 34,218,884 | 34,718,824 | Not significant | 1.33E-08 | Model failed | Model failed | Model failed |
| 17 | 39,422,031 | 39,459,777 | Not significant | 1.14E-07 | Model failed | Model failed | Model failed |

Supplementary Table 5 – Full list of haplotype blocks for which at least one RHM method significantly improved model fit for lamb jaw length. From left to right: chromosome, the haplotype block start (bp), haplotype block end (bp), and whether the inclusion of the regional GRM(s) significantly improved model fit (green), did not significantly improve model fit (yellow), or if a model failed to converge (red) for SNP-RHM, Hap-RHM, SNHap-RHM when compared to the null model, SNHap-RHM when compared to Hap-RHM, and SNHap-RHM when compared to SNP-RHM.
