## Supplementary Table 11 for "Evaluating regional heritability mapping methods for identifying QTLs in a wild population of Soay sheep"

| Trait | Chromosome | Haplotype number | SNP | SNP position | Method of identification |
|---|---|---|---|---|---|
| Birth weight | 1 | 3801 | oar3_OAR1_175180452 | 175180452 | GWAS |
| | 2 | 1589 | oar3_OAR2_81719138 | 81719138 | Conditional analysis |
| | 7 | 1173 | oar3_OAR7_54963456 | 54963456 | GWAS |
| Lamb foreleg | 16 | 1368 | oar3_OAR16_69873504 | 69873504 | GWAS |
| Lamb hindleg | 16 | 1368 | oar3_OAR16_69873504 | 69873504 | GWAS |
| Lamb metacarpal | 3 | 1713 | OAR3_100483326.1 | 94492563 | GWAS |
| | 16 | 1363 | s22142.1 | 69679810 | GWAS |
| | 19 | 952 | s74894.1 | 52470202 | GWAS |
| Adult August weight | 3 | 3696 | oar3_OAR3_209859215 | 209859215 | GWAS |
| | 6 | 630 | oar3_OAR6_35627634 | 35627634 | GWAS |
| | 9 | 847 | oar3_OAR9_36632898 | 36632898 | GWAS |
| Adult foreleg | 7 | 1918 | s48811.1 | 84579439 | GWAS |
| | 9 | 1165 | oar3_OAR9_50469115 | 50469115 | GWAS |
| | 11 | 668 | oar3_OAR11_30635038 | 30635038 | GWAS |
| | 16 | 1363 | s22142.1 | 69679810 | GWAS |
| | 19 | 950 | oar3_OAR19_52340442 | 52340442 | GWAS |
| Adult hindleg | 11 | 674 | oar3_OAR11_30832740 | 30832740 | GWAS |
| | 16 | 1363 | s22142.1 | 69679810 | GWAS |
| | 19 | 950 | oar3_OAR19_52340442 | 52340442 | GWAS |
| Adult metacarpal | 11 | 681 | oar3_OAR11_31212644 | 31212644 | GWAS |
| | 16 | 1393 | oar3_OAR16_71129444 | 71129444 | GWAS |
| | 17 | 1291 | oar3_OAR17_54802053 | 54802053 | GWAS |
| | 19 | 952 | oar3_OAR19_52459313 | 52459313 | GWAS |
| Adult jaw | 2 | *Falls between 2235 and 2236 | oar3_OAR2_137162126 | 137162126 | Conditional analysis |
| | 20 | 1031 | oar3_OAR20_46332617 | 46332617 | GWAS |

Supplementary Table 11 – Haplotype blocks containing the top GWAS SNPs from James et al. (2022). From left to right: trait, chromosome, haplotype block number, the SNP name, the SNP position, and whether the SNP was identified via GWAS or conditional analyses.
