## Supplementary Figure 1 for "Evaluating regional heritability mapping methods for identifying QTLs in a wild population of Soay sheep"

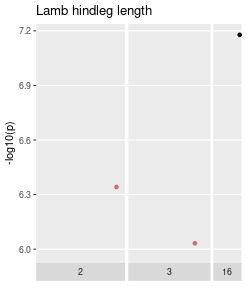

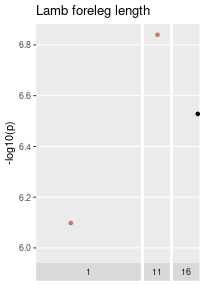

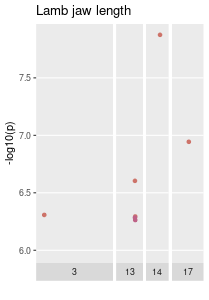

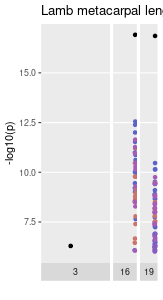

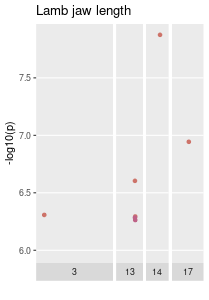

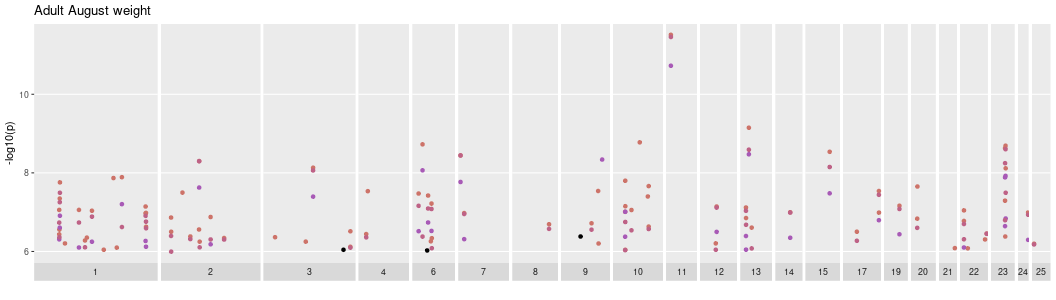

**A**

**E**

**B**

**C**

**D**

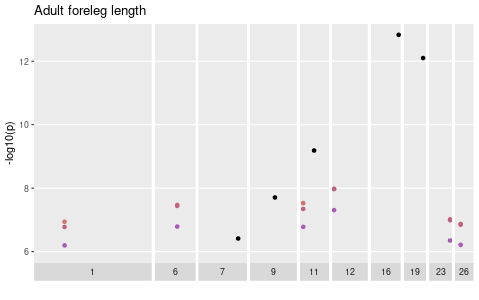

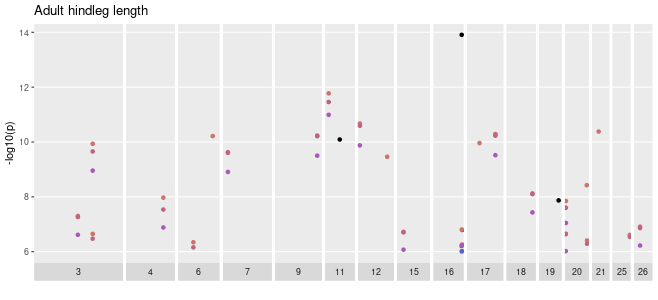

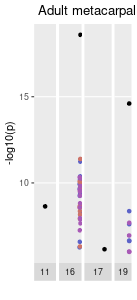

**F**

**G	H**

**Supplementary Figure 1 –** p values for haplotype block-trait associations for which at least one RHM model was significant for **A)** lamb foreleg length, **B)** lamb hindleg length, **C)** lamb metacarpal length, **D)** lamb jaw length, **E)** adult August weight, **F)** adult foreleg length, **G)** adult hindleg length, **H)** adult metacarpal length and **I)** adult jaw length. **J)** features the shared legend for these plots, indicating which colour represents each model. For traits that had significant SNP-trait associations in James et. al (2022), we included the p value of the top SNP for each association in black.

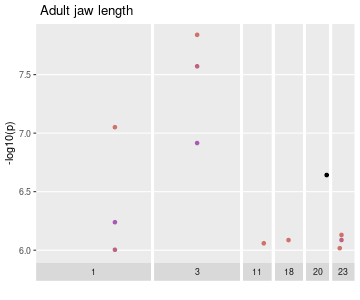

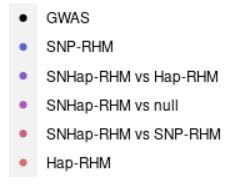

**I**

**J**
